# Targeted mutagenesis in *Ehrlichia canis* deleting the phage head-to-tail connector protein gene and its assessment as a vaccine candidate preventing canine ehrlichiosis

**DOI:** 10.64898/2026.01.29.702648

**Authors:** Dominica D. Ferm, Arathy Nair, Jonathan D. Ferm, Huitao Liu, Ying Wang, Liliana F. Crosby, Jodi McGill, Perle Latre De Late, Ian Stoll, Deepika Chauhan, Debika Choudhury, Swetha Madesh, Suhasini Ganta, Alexandra Burne, Sezayi Ozubek, Anish Yadav, Roman R. Ganta

## Abstract

*Ehrlichia canis* is primarily a *Rhipicephalus sanguineus* tick-borne rickettsial pathogen initially identified as causing canine monocytic ehrlichiosis, and infections in people have also been reported in Venezuela, Mexico, and parts of Europe. It is of high importance to have a vaccine suitable in protecting the canine host, which will aid in lessening *E. canis* infections also in people. Gene inactivation mutations in the phage head-to-tail connector protein genes (*phtcp*) from *E. chaffeensis* and *A. marginale* caused attenuated growth, and prior infection with the mutated bacteria induced protective immunity against wild-type bacterial infections in natural hosts, independent of blood-borne infection or tick-transmission infection. In the current study, we describe the development of targeted mutagenesis for the first time in *E. canis* genome and with a novel modification to avoid introducing antibiotic resistance cassettes to delete the *phtcp* ortholog from *E. canis*. The mutated *E. canis* was then assessed for its *in vivo* growth and the induction of host immunity exerted following the mutant infection aiding to protect against wild-type infection challenge in the canine host. We assessed systemic pathogen loads, hematological parameters, IgG immune responses, and plasma cytokines following the mutant infection relative to uninfected dogs. Similarly, the assessments were carried out following wild-type pathogen infections in dogs with or without prior mutant infection challenges. The study demonstrates that prior infection of dogs with the mutant induces immunity to prevent infection establishment by wild-type *E. canis.* Similarly, the mutant infection resulted in clear biological differences compared to the wild-type infection. This study establishes that the molecular genetic methods are broadly applicable to pathogens belonging to the family Anaplasmataceae and that the modified live vaccines with *phtcp* gene orthologs are valuable in reducing the diseases caused by the tick-borne rickettsial pathogens belong to Anaplasmataceae, including *E. canis*.

## 1. Introduction

Emerging and reemerging tick-borne rickettsial diseases remain a major public health concern in the USA and many parts of the world [1–5]. The diseases pose a significant risk to companion animal health, as well as contributing to global economic losses within the food animal production industry [6–9]. Despite some progress made, molecular genetic methods applicable to obligate intracellular pathogens remain a challenge [10–13]. Targeted gene inactivation methods are of great importance for investigations into defining the functional roles of genes [10, 14–16]. The availability of targeted mutagenesis methods greatly aids in advancing research in understanding the pathogenesis of obligate intracellular bacteria and extending research into the development of therapeutics and prevention methods [10, 12, 17–21].

Canine monocytic ehrlichiosis (CME), resulting from *Ehrlichia canis* infections, is the most prevalent tick-borne disease of dogs throughout the world, including in the USA [22–26]. The primary transmitting vector is *Rhipicephalus sanguineus sensu lato* (s.l.) complex which comprises of multiple, closely related, and morphologically similar species which are commonly known as the brown dog tick [27–29]. While the tick complex prefers dogs as the primary host and resides indoors, it also feeds on people [27, 28]. Human infections with *E. canis* are reported from several Western hemisphere and European countries, such as Venezuela, Mexico, Costa Rica, Italy, and Montenegro [30–35]. *E. canis* tropism is to the immune cells, monocytes, and macrophages, leading to a multisystemic disease ranging from acute to subclinical and chronic phases [24, 36, 37]. CME is characterized by nonspecific clinical signs such as fever, lethargy, inappetence, and several hematological abnormalities, including thrombocytopenia, leukopenia, and anemia [38]. Most immunocompetent dogs will naturally clear the clinical signs without any treatment. However, the disease can potentially progress to a chronic phase, causing aplastic anemia or severe bone marrow aplasia, known as myelosuppression, developing into life-threatening pancytopenia [39–41]. If left untreated, death can occur, resulting from septicemia or other complications related to the disease [39–41]. Considering the impact of *E. canis* infections on dogs and humans and the limited treatment options, mainly the use of doxycycline [42–46], it is of critical importance to have a vaccine that protects the canine host [37], which will also aid in reducing the risk of human infections with the pathogen. Despite a prior study reporting that a cell culture-derived attenuated strain provided immune protection against virulent challenge [36], to date, there are no follow-up investigations to advance vaccine research in preventing CME. Additionally, this study did not describe how bacterial attenuation is achieved at the genomic level [36].

We previously developed a homologous recombination-based molecular genetic system in *Ehrlichia chaffeensis* [11, 47], a member of the Anaplasmataceae family of pathogens and responsible for causing infections in people and several other vertebrates, including dogs [2, 22, 48–53]. The mutational methods have been valuable in creating mutations in several bacterial genes, including disrupting and restoring a gene function [11]. We also reported that inactivation of the gene encoding a membrane-bound phage head-to-tail connector (phtcp) of *E. chaffeensis* causes attenuated growth *in vivo* in both reservoir (white-tailed deer) and incidental (dog) hosts [17, 47](naturally known to acquire infections from ticks, while inducing a long-lasting immunity conferring protection against wild-type infection challenge by both intravenous (IV) inoculation and tick transmission [17, 18, 20, 21, 47]. Further, we reported that the *E. chaffeensis phtcp* gene inactivation results in the pathogen’s inability to obtain metal ions to support its growth in infected host macrophages [54]. Since the *phtcp* gene orthologs are well-conserved in all known *Anaplasma* and *Ehrlichia* species [54], we extended similar targeted mutagenesis development by deleting the *phtcp* ortholog from another related *Anaplasmataceae* pathogen, *A. marginale* [12]. Furthermore, we reported that vaccination of cattle with the genetically modified *A. marginale* mutant lacking the *phtcp* displays significantly attenuated growth and induces immunity to prevent severe clinical bovine anaplasmosis resulting from IV infection or from infected ticks [12, 19].

In the current study, we developed the first targeted deletion mutation in *E. canis* while not introducing any antibiotic resistance genes as part of the mutational strategy. With no vaccines available to prevent CME, we assessed whether the *phtcp* deletion mutant, as a modified live attenuated vaccine (MLAV), can serve as a vaccine candidate to support improving companion animal health (supplementary Fig S1). Vaccination of dogs with the *E. canis phtcp* gene deletion mutant resulted in the canine host inducing immunity that is sufficient in preventing wild-type pathogen infection establishment.

## 2. Materials and methods

### 2.1 Culturing E. canis

Wild-type and the genetically modified mutant *E. canis* Jake isolate were cultivated in *Ixodes scapularis* cell line culture (ISE6) (ATCC# CRL-3576) at 34°C in the absence of CO_2_ as described earlier [55]. Wild-type and mutant *E. canis* were also cultivated in the canine macrophage cell line, DH82 (ATCC# CRL-10389), at 37°C in an incubator with 5% CO_2_.

### 2.2 Generation of Ecaj_0381 deletion plasmid construct

Homologous arms of 0.84 kb and 0.86 kb in length spanning the 5’ and 3’ to the targeted deletion region of *Ecaj_0381* gene (the *phtcp* gene homolog) of *E. canis* (GenBank # CP000107) were amplified by PCR using specific primers (Table S1) and using wild-type bacterial genomic DNA as the template. The amplicons were cloned upstream and downstream to the previously described pGGA plasmid vector (New England Biolabs, Ipswich, USA) containing the *A. marginale amtr* promoter to drive the expression of the mCherry gene coding sequence and *aadA* gene encoding resistance to streptomycin and spectinomycin (pLox cisA7Himar Ch-SS) [56]. A Golden Gate Assembly kit (New England Biolabs, Lpswich, MA, USA) was used to reassemble the fragments in the following order: 5’ *E. canis* homology arm, *Amtr-*mCherry-aadA segment, and 3’ *E. canis* homology arm. The 5’-3’ homologs are referred to here as the Left homologous arm (LHA) and the Right homologous arm (RHA). The recombinant plasmid was transformed into the DH5α strain of *E. coli* and used to generate plasmid DNA by following the standard molecular biology protocols [57]. The proper assembly of the recombinant plasmid was confirmed by DNA sequence analysis using T7 and SP6 promoter primers (Integrated DNA Technologies, Coralville, IA, USA) having binding sites upstream and downstream to the insertion segment within the pGGA plasmid backbone. Subsequently, the *aadA* gene was deleted from the recombinant plasmid using the Q5 Site-Directed Mutagenesis Kit (New England Biolabs, Ipswich, USA) according to the manufacturer’s protocol, resulting in a new recombinant construct designated pGGA-Ecaj_0381-KO-Amtr-mCh. This modified plasmid was then used as the template to amplify the insertion segment containing the LHA, the *Amtr-*mCherry segment, and the RHA using the primers (RRG2015 and RRG2048) listed in Table S1. The PCR amplicons were then purified using a QIAquick PCR Purification Kit (Qiagen, Germantown, MD) as in [11].

### 2.3 Generation, clonal purification, verification, and propagation of E. canis phtcp mutant

The *E. canis phtcp* deletion mutation was generated as per prior reported methods [11, 12] with a modification to exclude the use of an antibiotic selection marker. The cultures were maintained in the media without added antibiotics, and the media were changed once a week for the first three weeks and twice a week thereafter. Cultures were assessed for the mutant expressing mCherry by fluorescent microscopy. To achieve clonal purity of the mutant from wild-type *E. canis,* mutant bacteria were enriched through repeated serial dilutions of cultured bacteria, which took several months. When approximately 10% of the culture was expressing mCherry, the media supernatant from infected cultures containing cell-free bacteria was collected and used to infect a monolayer of ISE6 tick cells in six-well plates. The day after infecting the monolayer, all media was removed and replaced with 2 ml of a modified infection medium containing 0.4% low-melting-temperature agarose, which created a thin agarose overlay allowing the development of a plaque of cells expressing mCherry. After the agarose had set overnight, 4 mL of culture media was added to the agarose layer, and cultures were monitored for 4 weeks. When colonies of mCherry expressing fluorescent bacterial colonies were visible (between 3 and 4 weeks), liquid media was removed, and a pipette tip was used to carefully recover mCherry expressing fluorescent cells from beneath the agarose layer. The recovered cells were then diluted in 0.5 ml culture media in sterile 1.5 ml microcentrifuge tubes, pipetted to disrupt cell clumps and agarose, and added to wells of a fresh 6-well plate having an ISE6 cell monolayer maintained in infection media. A total of three sequential passages were performed to isolate the purified mutant. Presence of the mutant expressing mCherry was monitored and confirmed by fluorescence microscopy, and images were captured using BioTek Cytation C10 Confocal Imaging Reader (Agilent, USA). The clonal purity of the mutant was independently established by PCR, Southern blot analysis, and whole-genome sequencing analysis as outlined below.

A PCR assay was performed using genomic DNA recovered from the mutant to define the clonal purity using the primers (RRG2015 and RRG2048) listed in Table S1. The primers were designed to target the genomic regions upstream and downstream of the gene deletion region (listed in Table S1). The PCR assays were performed in 25 μl reactions in 1x Q5 reaction buffer containing 2 mM MgCl_2_, 0.5 mM of each dNTP, 0.2 μM of each of the primers, 1 unit of Q5 high-fidelity DNA polymerase (New England Biolabs, Ipswich, MA, USA), and genomic DNA from wild-type or mutant organisms as the templates. The PCR cycling conditions for the first two PCRs were 98°C for 30 s, followed by 35 cycles of 98°C for 10 s, 65°C for 30 s, and 72°C for 2 min 30 s, then 72°C for 3 min, and a final hold at 4°C. The presence of a specific PCR amplicon was confirmed by resolving DNAs on a 1 % agarose gel containing ethidium bromide and visualizing it under a UV transilluminator. To independently confirm the clonal purity of the mutant, Southern blot analysis was performed using the mutant culture-derived genomic DNA as well as the wild-type *E. canis* genomic DNA; the DNAs were digested with BamH1, Sph1, or a combination of BamH1and Sph1 restriction enzymes by following the standard molecular biology protocols [57]. The digested DNAs were resolved on a 0.9 % agarose gel, transferred to a nylon membrane, and subjected to blot analysis using a DNA segment to be retained in the restriction-digested DNA fragments surrounding the genetically modified region spanning DNA from the LHA or insertion-specific mCherry gene segment DNA as a probe. Whole-genome sequencing analysis was performed to further confirm the clonal purity of the mutant. Genomic DNA extracted from the mutant-infected DH82 was sequenced using a hybrid sequencing approach that combined the Oxford Nanopore MinION and Illumina NextSeq 2000 platforms (Plasmidsaurus Inc., Kentucky, USA). To remove Nanopore adapters and to perform quality filtering, we used Porechop v.024 [58] and Filtlong v.0.2.1 (https://github.com/rrwick/Filtlong) with default settings. The remaining high-quality reads were mapped to the *E. canis* Jake isolate reference genome (CP000107) and for the presence of *Ecaj_038*1deletion construct sequences including the *Amtr* promoter and *mCherry* using Minimap2 v.2.28 [59]. Mapped reads were extracted via samtools v.1.21 [60] and assembled *de novo* using Flye v.2.9.6 [61]. To improve the assembly, the Illumina NextSeq2000 reads were groomed and filtered for quality using Trimmomatic v.0.39 [62]. The high-quality Illumina reads were then mapped to the draft assembly using Bowtie2 v.2.5.3 [63], followed by polishing using Polypolish v.0.6.0 [64].

RNA analysis was performed using total RNA isolated from the mutant and the wild-type *E. canis*, as described elsewhere [65], to assess mRNA expression from the gene *Ecaj_0381* and from a gene upstream and downstream to the mutant site; *Ecaj_0383* and *Ecaj_0380*, respectively. After verifying the removal of any residual genomic DNA from the RNAs by DNase I treatment using Q1 RNase-Free DNase (Promega, Madison, WI, USA) per the manufacturer’s recommendation, first-stand cDNAs were synthesized using SuperScript III First-Strand Synthesis System for RT-PCR kit (Invitrogen, Carlsbad, CA, USA) and used as templates to perform RT-PCR analysis. Primers for the three gene targets were listed in Table S1. Polymerization was performed with 2 μl of cDNA template using GoTaq Green DNA polymerase per the manufacturer’s instructions (Promega, Madison, WI, USA). The PCR products were resolved in a 1.5% agarose gel stained with ethidium bromide, and a UV transilluminator was used for visualization to identify the target PCR products.

### 2.4 Infection experiments in dogs with the mutant E. canis

Infection experiments in dogs of both sexes were performed. The study design was reviewed and approved by the University of Missouri’s Animal Care and Use Committee (UM ACUC) and executed as per the approved protocol and as per compliance with the Public Health Service (PHS) policy on the humane care and use of laboratory animals and the U.S. Department of Agriculture’s Animal Welfare Act and Animal Welfare Regulations (https://olaw.nih.gov/policies-laws/phs-policy.htm). We obtained purpose-bred laboratory-reared beagle dogs of both sexes about 8 months of age from a USDA approved commercial vendor (Ridglan Farms, Inc. Mt. Horeb, WI). All dogs were allowed to acclimate for two weeks. During this period, 4 ml each of blood samples in EDTA tubes were collected from all dogs and used to confirm that none of the animals had current or past *E. canis* infections. The analysis was performed by subjecting total DNAs recovered from blood by pathogen-specific PCR. Additionally, plasma samples recovered from the bloods were evaluated by ELISA analysis for the pathogen-specific antibodies by ELISA. All dogs tested negative by PCR and ELISA. The infections were performed using either the *in vitro* cultured mutant organisms or the wild-type *E. canis* cultured in DH82 cells. Before mutant or wild-type infection, animals were administered Benadryl orally at 2 mg/kg approximately 30 min prior to the IV infection challenge. For mutant *E. canis* infection experiments, dogs were administered about 1.25 × 10^8^ culture-derived mutant organisms re-suspended in 1 ml of 1 x PBS (n=5) intravenously (Group 1). Similarly, a group of dogs (n=4) that did not receive the mutant were injected with 1 x PBS only and served as controls (Group 2). Both groups of dogs were then challenged with wild-type *E. canis* on day 29 following the mutant infection by IV using 2 ml each of cultured organisms (about 1.25 x 10^8^ bacteria per dog). All dogs were monitored for signs of infection, changes in body temperature, weight, and Complete Blood Count (CBC) analysis. Blood samples in EDTA tubes were collected over time and used to define the presence of infection by *in vitro* culture recovery and PCR analysis as described in the previous section. Further, total RNAs were isolated from 250 µl of each blood sample at each time point throughout the study period. Quantitative reverse transcriptase polymerase chain reaction (qRT-PCR) assays targeting the bacterial 16S rRNA (using primers listed in Table S1) were performed as described in our previous work to further define the infection status [66]. After day 72 post-challenge, all animals received doxycycline at a dose of 10 mg/kg body weight once a day orally for 28 days as per Merck veterinary manual. Blood samples collected on days 11 and 32 post-doxycycline treatment were assessed by PCR using primers RRG2015 and RRG2048, listed in Table S1.

### 2.5 qRT-PCR

Two microliters of total RNA recovered from 250 μl of blood samples collected in EDTA tubes from dogs at various time points were assessed by qRT-PCR to detect and quantify *E. canis* RNA. This was performed according to the manufacturer’s instructions using the SuperScript III Platinum One-Step RT-qPCR Kit (Thermo Fisher Scientific, USA). The qRT-PCR conditions were followed as previously described [66].

### 2.6 Assessment of IgG response by ELISA

Whole-cell antigen was prepared and used to perform an enzyme-linked immunosorbent assay (ELISA) as described earlier [17]. Briefly, ninety-six-well Immulon™ 2HB microtiter plates (Thermo Fisher Scientific, Waltham, MA) were coated with 2 μg/ml *E. canis* WCA in the ELISA coating buffer and incubated overnight. Plates were then blocked with 1% BSA for one hour at 37°C, followed by 2-3 hours of incubation at 37°C with 1:200 diluted plasma collected at different time points post-mutant infection, or following infection challenges with wild-type *E. canis*. The plates were washed three times with 1X PBS containing 0.05% Tween 20 (1X PBST), then incubated with 100 μl of horseradish peroxidase (HRP)-conjugated anti-canine total IgG at a 1:50,000 dilution (PA1-29738, Thermo Fisher Scientific, USA). The plates were washed three times, then incubated with TBM substrate for about 5-8 min at room temperature. The reactions were stopped with the stop buffer having 1 M phosphoric acid, and the absorbance at 450 nm was measured using a Tecan Infinite M Nano (Männedorf, Switzerland).

### 2.7 Plasma cytokines analysis

The commercially available Cytokine/Chemokine/Growth Factor 11-Plex Canine Procarta Plex™ Panel 1 kit (Thermo Fisher Scientific, USA) was used to measure secreted cytokines and chemokines using Luminex xMAP technology. The assays were performed according to the manufacturer’s instructions using plasma samples collected in EDTA tubes on the day before mutant infection and on days 3, 7, and 14 post-immunization/infection. Samples were also taken after *E. canis* wild-type infection challenges for the same time points from all groups of dogs.

### 2.8 Preparation of peripheral blood mononuclear cells (PBMC) to perform ELISPOT and ELISA assays for measuring the IFN-γ expression

Blood samples were collected from dogs at different time points in 8 ml sodium citrate CPT tubes (BD Biosciences, San Jose, CA, USA) on the days preceding any infection or vaccination, and then on days 7 and 14 post-mutant infection challenges, and after wild-type infection challenges. The samples were centrifuged at 1500 x g for 30 min at room temperature, and the separated cells were thoroughly mixed before being shipped overnight to Iowa State University in cold packs. The harvested PBMCs were washed twice with cRPMI media, followed by cell counting and viability assessment. The cells were then adjusted to 4 × 10^6^ PBMCs/mL, and ELISPOT and ELISA assays were performed to measure IFN-γ expression by stimulating PBMCs with *E. canis* whole-cell antigens, as described in [20].

### 2.9 Statistical analysis

Using GraphPad Software (GraphPad Software, La Jolla, CA, USA), the one- and two-way ANOVA tests with multiple comparisons, along with t-tests, were performed to identify statistical differences among groups of dogs following mutant infections and following wild-type infection challenges. As we found no evidence of sex-specific differences for all parameters assessed, the combined data were presented and described in the study.

## 3. Results

### 3.1 Preparation of the homologous recombination construct for use in deleting the phtcp gene from E. canis

To generate the targeted deletion mutation of the *Ecaj_0381* gene encoding for phtcp from the *E. canis* strain genome (Jake isolate) (GenBank # CP000107), the homology arms spanning the genomic regions upstream and downstream to the *phtcp* gene, respectively, were cloned into a plasmid construct (pGGA). *A. marginale tr* gene promoter (*Amtr*) and the mCherry reporter gene coding sequence were inserted between the left and right homology arm segments (LHA and RHA) to allow mCherry expression in the mutant. The resulting recombinant plasmid construct, referred to as pGGA-Ecaj_0381-KO-Amtr*-*mCh, lacks any antibiotic selection markers while the mCherry coding sequence facilitates driving its expression by the *Amtr* promoter to track the mutant and its isolation (Fig 1A). The plasmid was then used as a template to generate homologous recombination segments by PCR to generate the construct segment, which contained both the 5’ and 3’ homology arms and the *Amtr*-mCherry segment. Genomic coordinates of the homology arms, position of restriction enzyme sites, and the *Amtr*-mCherry segments are detailed in (Fig 1B).

**Figure 1.**
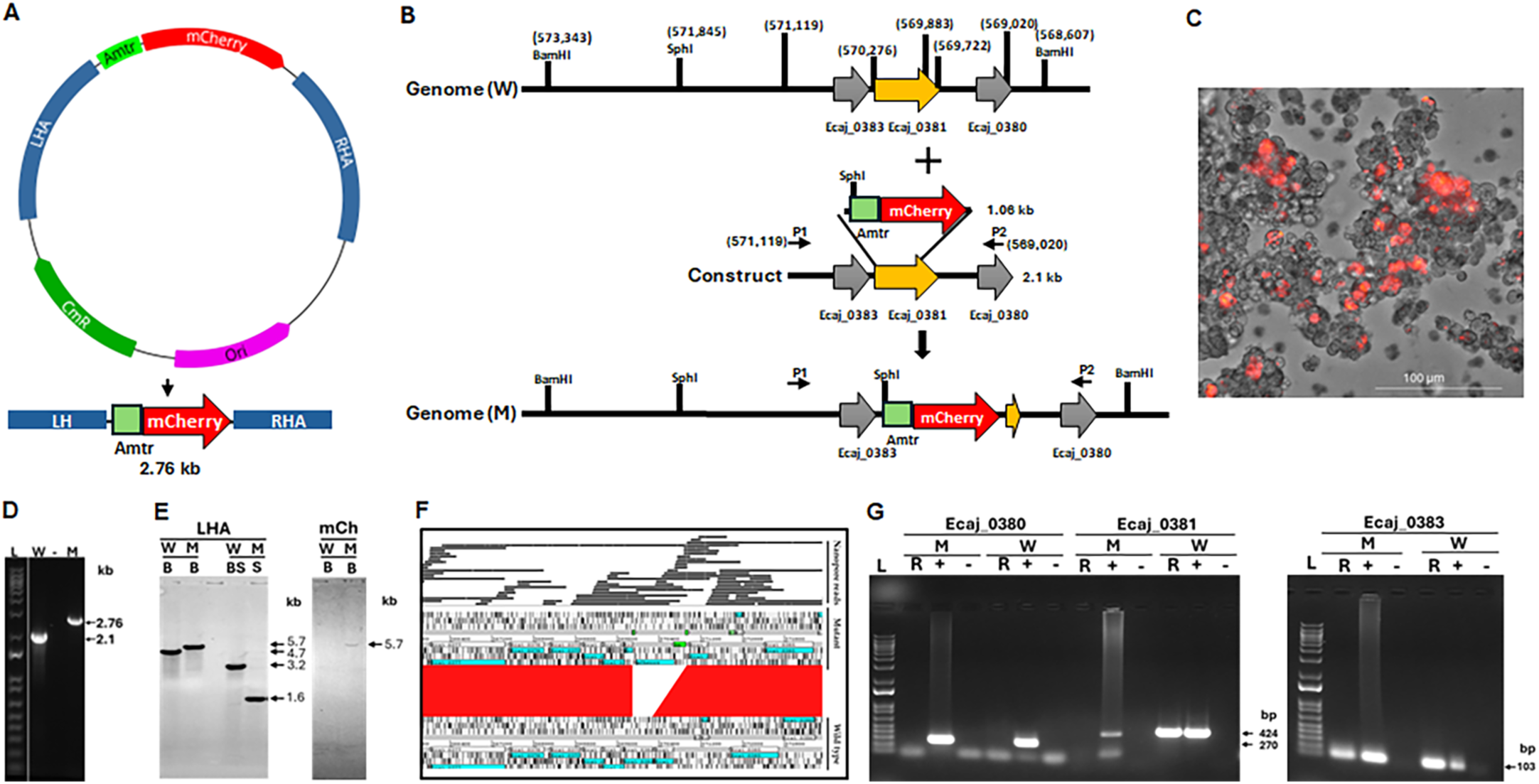
The development of Ecaj_0381 (*phtcp*) gene deletion mutation from the *E. canis* genome. A) Plasmid map of pGGA-Eca_0381-KO-Amtr-mCh A. The plasmid was used to amplify the left (LHA) and right (RHA) homology arms flanked the *Amtr* promoter to drive expression of the mCherry reporter gene segment. B) Schematic representation of the *E. canis* genomic region targeted mutagenesis generation from wild-type (W) to mutant (M) by allelic exchange. The genomic coordinates of the LHA and RHA, restriction enzyme sites used for mutation verification (BamHI and SphI), and the size of the inserted *Amtr*-mCherry fragment are shown. Black arrows indicate the primers binding sites; P1 (RRG2051) and P2 (RRG2048) used to confirm *Ecaj_0381* gene deletion. C) The gene deletion mutant growth in ISE6 tick cell culture expressing mCherry (confirmed by confocal microscopy using 40x magnification lenses at 400 x magnification. D) PCR analysis confirming the mutation. Primers annealing upstream and downstream of the mutation insertion region were amplified and resolved on an agarose gel which yielded the expected larger and smaller products from the mutant (M) and wild-type (W) DNAs, respectively (L, 1 kb plus DNA ladder). E) Southern blot analysis of W and M genomic DNAs digested with BamHI or double digested with BamHI and SphI. The digested genomic DNAs resolved on an agarose gel and transferred to a nylon membrane were hybridized with probes specific to the LHA or mCherry gene coding sequences, which verified the presence of the predicted segments in the clonally purified mutant DNA and for wild-type DNA. F) Nanopore sequencing reads used to assemble the genome of Ecaj_0381 mutant compared with the *E. canis* str. Jake reference genome (CP000107). The genome alignments revealed the expected Ecaj_0381 gene deletion in the mutant and inserted fragment in its place. Sequences flanking the transposon insertion site shared 99% identity. G) The impact of *E. canis Ecaj_0381* (*phtcp)* gene deletion was assessed for the gene expression from it and two genes located upstream and downstream of the deleted gene. Transcriptional analysis of RNA recovered from the mutant and wild-type *E. canis* was assessed by RT-PCR targeting *Ecaj_0380*, *Ecaj_0381*, and *Ecaj_0382*. The assay included wild-type genomic DNA as the template for the PCR to serve as the positive control (+), no template PCR as the negative control (-), and cDNA as the template to represent RNA samples (R) from the organisms. (L, 1 kb plus DNA markers). The mutant cDNA lacked the expected amplicons only for the *Ecaj_0381*, but not in the cDNA from wild-type *E. canis,* while RNA expression from the *Ecaj_0380* and *Ecaj_0383* genes was similar for both cDNAs.

The targeted mutagenesis protocol for *E. canis phtcp* gene deletion was followed as per previously reported methodology for *E. chaffeensis* and *A. marginale* [11, 12]. The method involved utilizing 20 µg of linear DNA fragments for electroporation into purified *E. canis* organisms derived from ISE6 cells (*Ixodes scapularis* embryonic cells) cultures. The presence of *E. canis* mutant organisms expressing mCherry fluorescent protein in ISE6 cells was monitored; the mutant cultures expressing mCherry appeared after three weeks post-electroporation experiment (Fig 1C). Clonal purification of the mutant was achieved by enrichment of mutant bacteria by serial dilution through several passages of mutant and wild-type infected tick cells, followed by a 0.4% low melt agarose layered on top of the tick cell monolayer to facilitate the recovery of purified mutant colonies expressing mCherry; three sequential passages were performed to isolate the purified mutant. (More details of the method were included in the methods section). The presence of a clonally pure mutant was confirmed by three independent molecular methods: PCR analysis to identify the presence of a predicted larger amplicon in the mutant compared to the wild-type (Fig 1D), Southern blot analysis following restriction enzyme digestions with BamHI (B) or SphI (S) or double digestions with both enzymes (BS) utilizing either a DNA probe specific to the LHA or the mCherry coding sequence (mCh) (Fig 1E), and by whole genome sequencing analysis (Fig 1F). Isolated genomic DNA of the mutant *E. canis* included only the predicted 2.7 kb mutant-specific amplicon contrary to 2 kb observed for the wild-type genomic DNA, indicating the absence of wild-type genomic DNA (Fig 1D). Similarly, Southern blot analysis with the LHA genomic probe identified predicted restriction fragments of 5.7 kb for BamHI or 1.6 kb with SphI digested genomic DNA from the mutant, while the predicted 4.7 kb with BamHI and 3.2 kb with BamHI and SphI double digestion DNA fragments were evident when wild-type genomic DNA was used (Fig 1E, left panel). The anticipated 5.7 kb size for the BamHI-digested DNA was observed only in the mutant when the mCherry DNA probe was used demonstrating that there were no other locations in the genome where the insertion mutations resulted, and it was absent in wild-type DNA, similarly digested (Fig 1E, right panel). Further, whole genome sequence analysis of the mutant revealed the presence of the inserted sequence of *Amtr* promoter followed by mCherry coding sequence and the absence of the *Ecaj_0381* gene in the mutant (Fig 1F). Further, the whole genome sequence analysis confirmed the presence of mCherry sequences only at the targeted mutational site. We then performed RNA analysis by reverse transcriptase PCR (RT-PCR) which revealed the anticipated absence of transcript for gene *Ecaj_0381* in the mutant but not in the wild-type, while the RNA expression from genes upstream and downstream (*Ecaj_0383* and *Ecaj_0380*) remained similar for the mutant and wild-type (Fig 1G).

### 3.2 Impact of the phtcp gene deletion on the progression of E. canis infection in a physiologically relevant host

The canine infection model study flow chart is included (supplementary Fig S1). We assessed the impact of the *phtcp* gene deletion on the pathogen’s *in vivo* growth in the canine host, as it acquires the pathogen naturally. Five beagles (Group 1) were injected IV with approximately 1.25 x 10^8^ mutant organisms per dog and monitored for the systemic infection-associated changes twice a week for four weeks. The assessment included evaluating clinical symptoms, changes in CBC, presence of the mutant in circulation by PCR analysis, plasma cytokine secretion, and the pathogen-specific IgG response. Four beagles (Group 2) were included as uninfected controls. Over the course of the study, none of the mutant-infected or uninfected control beagles developed clinical signs such as fever or weight changes. Mutant genomic DNA was not detected in any of the controls (Group 2) but was detected in the blood samples of all five mutant-inoculated beagles (Group 1) by mutation region-specific PCR analysis (Table 1). Blood-derived RNA was similarly assessed by qRT-PCR targeting the *E. canis* 16S rRNA, and RNA copies were estimated as per our previously described method [66]. We detected the presence of bacterial RNA with a sharp increase over time, reaching its highest level on day 14 post-infection, followed by a marked decline thereafter (Fig 2 and supplementary Table S1). Since *E. canis* infections in dogs are known to alter hematological parameters, we performed CBC analysis, which revealed significant changes in the blood profiles of mutant-infected dogs compared to uninfected dogs (Fig 3). The mutant infection-specific changes included a substantial decline in the platelets (Fig 3A), red blood cells (RBC) (Fig 3B), and white blood cells (WBC) (Fig 3C). The mutant-infected animals developed an IgG response against the *E. canis* antigens, as evidenced from day 7 post-infection, while no IgG response was detected in uninfected dogs (Fig 4).

**Figure 2.**
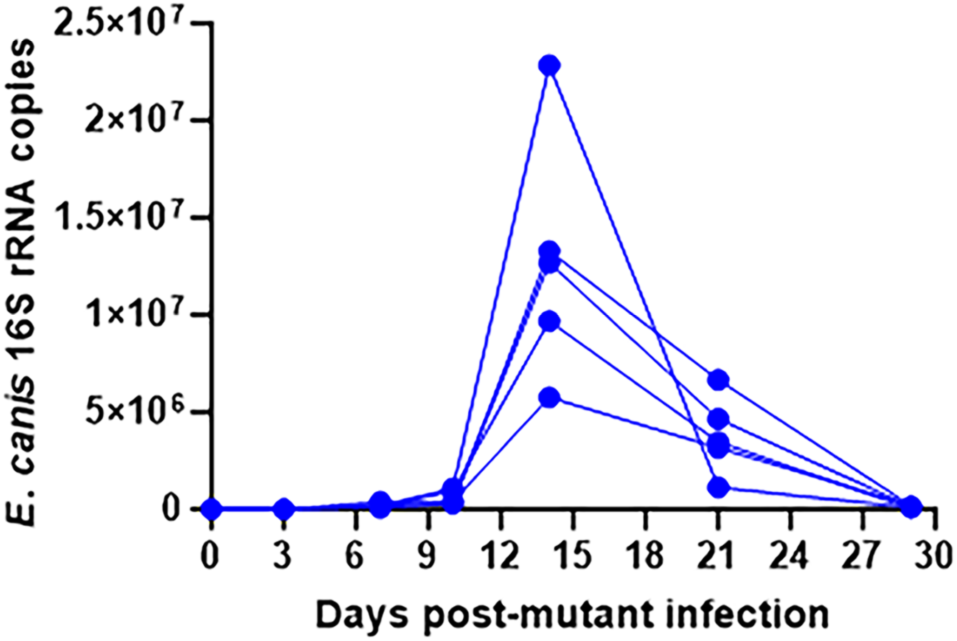
*E. canis 16s rRNA* copy numbers in blood-derived RNA assessed by TaqMan qRT-PCR following infection with the *phtcp* mutant. Total RNA was isolated from blood samples collected at different time points following mutant infection in the canine host (n = 5). The qRT-PCR analysis was performed as per our previous study [66] to quantify *E. canis* small subunit ribosomal RNA copies per 25 μl of blood.

**Figure 3.**
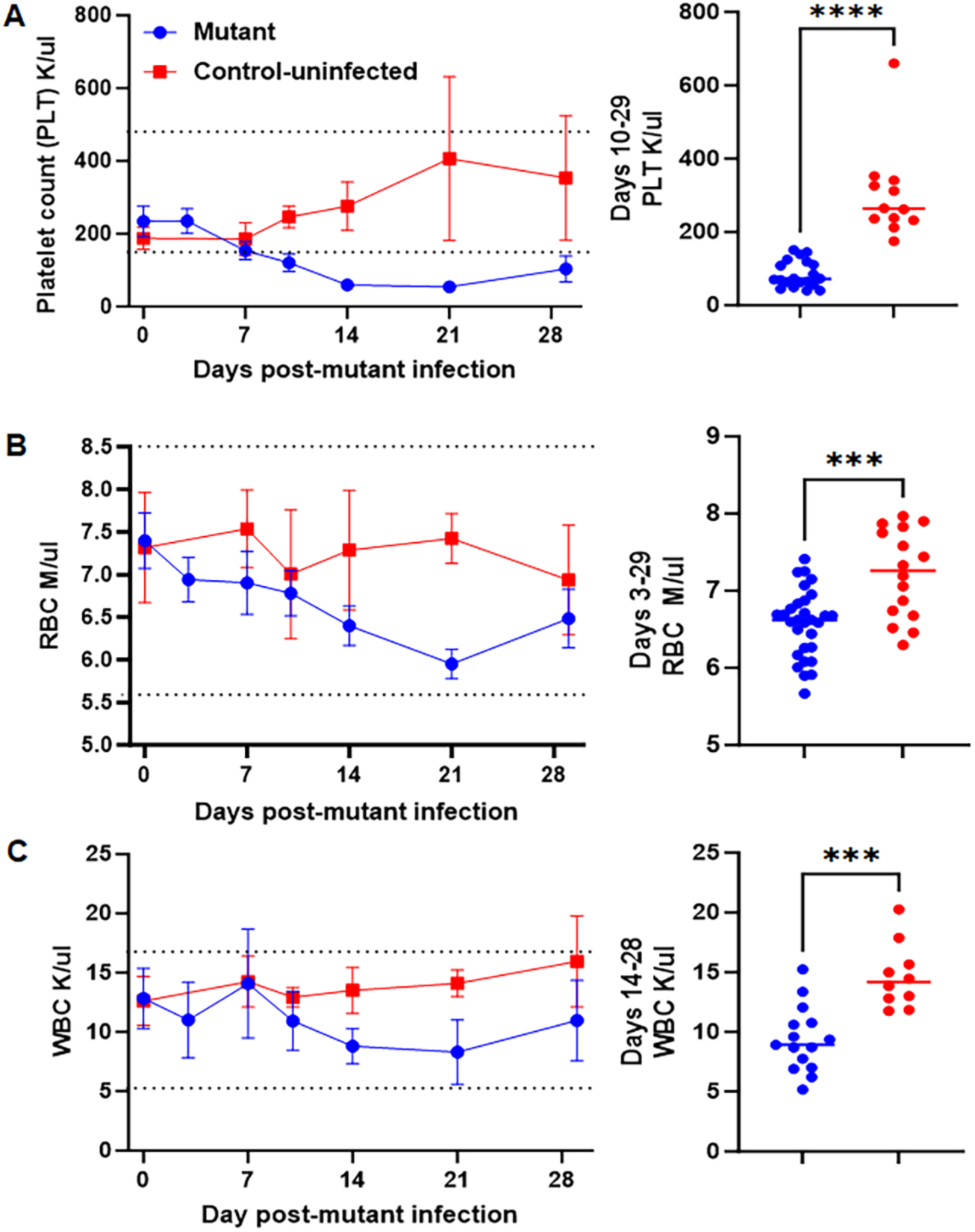
The complete blood count (CBC) assessment following the mutant *E. canis* infection in the canine host. The CBC analysis performed over time following mutant *E. canis* infection revealed a significant drop in the platelet counts (P < 0.0001 ****) (panel A), RBC counts (P = 0.0007 ***) (panel B), and WBC counts (P < 0.0001 ****) (panel C), compared to uninfected dogs. The dotted lines indicate normal canine reference values, as per the Merck Veterinary manual. Statistical significance was measured using an unpaired t-test with Welch’s correction. Statistical significance was calculated from day 10 to 29, when values started to decline following mutant infection, and compared with the uninfected control group (right panels).

**Figure 4.**
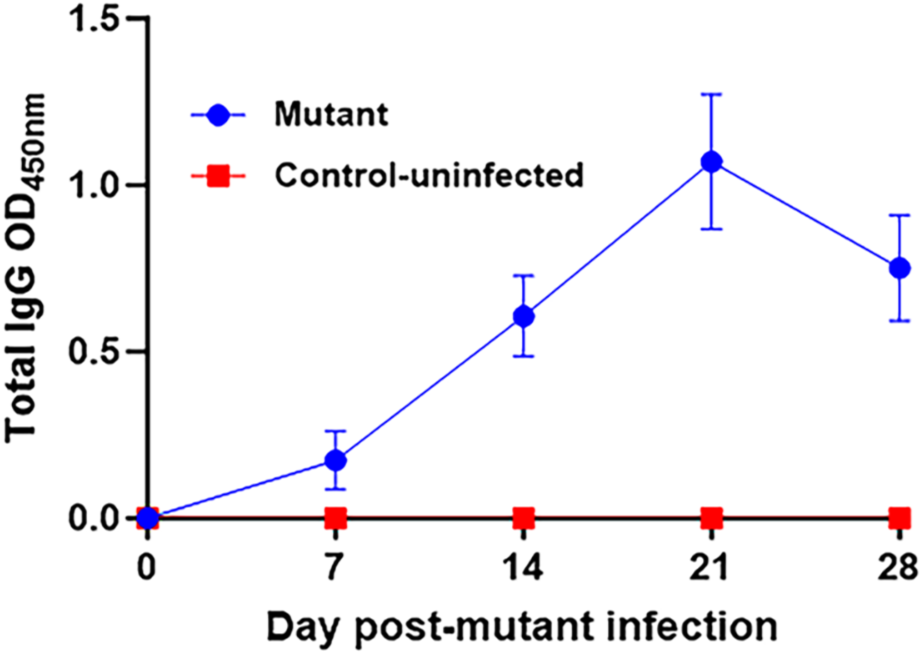
Total IgG response to *E. canis* antigens in dogs was measured by ELISA using plasma samples collected weekly following the mutant infection. Specific antibody responses were observed from day 7 post-infection in the mutant-infected group of dogs but not in the uninfected control dogs.

**Table 1.** *E. canis* mutant infection assessment in the canine host (Mutant-specific PCR).

|  | Dog ID | Days post mutant infection |  |  |  |  |  |  |
| --- | --- | --- | --- | --- | --- | --- | --- | --- |
|  |  | 0 | 3 | 7 | 10 | 14 | 21 | 29 |
| Mutant infected dogs (Group 1) | AHL (M) | - | - | + | + | + | + | - |
|  | BAK (F) | - | - | + | + | + | + | - |
|  | BGK (F) | - | - | + | + | + | + | + |
|  | CUL (M) | - | - | + | + | + | + | + |
|  | DWL (M) | - | - | + | + | + | + | - |
| Dogs that did not receive the mutant infection (Group 2) | AVK (F) | - | - | - | - | - | - | - |
|  | BCK (F) | - | - | - | - | - | - | - |
|  | ECL (M) | - | - | - | - | - | - | - |
|  | GVL (M) | n/a | n/a | n/a | n/a | n/a | - | - |

\* + and – refer to PCR positives and negatives, respectively, and n/a refers to samples not available. M and F refer to male and female dogs, respectively.

### 3.3 Evaluation of the E. canis phtcp gene deletion mutant as a vaccine

*We* then evaluated if the prior injection with the *E. canis phtcp* gene deletion mutant serves as an MLAV in offering protection against wild-type *E. canis* infection challenge. Approximately 1.25 x 10^8^ cultured *E. canis* wild-type bacteria were IV administered into all five Group 1 beagles on day 29 post-mutant infection. Similarly, the control group beagles (Group 2) received the wild-type *E. canis* to serve as the infection controls.

#### 3.3.1 Prior mutant infection offered protection against Wild-type E. canis

The mutant-infected group following wild-type infection challenge remained positive for the mutant-specific PCR throughout the 72-day assessment period when assessed using blood-derived DNAs, although the number of dogs testing positive declined with time. The Group 2 controls tested negative for the mutant-specific DNA (Table 2). All dogs in the MLAV group remained negative for wild-type *E. canis* throughout the 72-day monitoring period following the wild-type infection challenge.

**Table 2.** *E. canis* infection assessment by PCR targeting the presence of the mutant segment spanning the mutated region (Mutant-specific PCR).

|  | Dog ID | Days post-wildtype infection challenge* |  |  |  |  |  |  |  |  |  | Post-Doxycycline** |  |
| --- | --- | --- | --- | --- | --- | --- | --- | --- | --- | --- | --- | --- | --- |
|  |  | 3 | 7 | 10 | 14 | 18 | 21 | 28 | 35 | 49 | 72 | 11 | 32 |
| Mutant-infected followed by wild-type infection challenge (Group 1) | AHL (M) | + | - | - | + | + | - | + | - | - | - | - | - |
|  | BAK (F) | + | - | + | + | + | + | + | + | - | - | - | - |
|  | BGK (F) | - | - | - | + | + | + | + | + | - | - | - | - |
|  | CUL (M) | + | - | - | - | + | - | - | - | - | + | - | - |
|  | DWL (M) | + | - | + | + | + | - | + | - | + | - | - | - |
| Dogs receiving only wild-type infection (Group 2) | AVK (F) | - | - | - | - | - | - | - | - | - | - | - | - |
|  | BCK (F) | - | - | - | - | - | - | - | - | - | - | - | - |
|  | ECL (M) | - | - | - | - | - | - | - | - | - | - | - | - |
|  | GVL (M) | - | - | - | - | - | - | - | - | - | - | - | - |

\* + and – refer to PCR positives and negatives, respectively. M and F refer to male and female dogs, respectively.
\*\*All dogs in these two groups were treated with doxycycline for a month after 72 days of study assessment and then assessed for the presence of infection by PCR at the two time points listed.

This was consistent regardless of the detection methods used: culture recovery, which was performed on days 14 and 21, or by a conventional PCR assay (Table 3). In contrast, all dogs in the infection control group (Group 2) challenged with wild-type *E. canis* tested positive by both PCR beginning on day 10 post-challenge and by culture recovery assessment (assessed only on days 14 and 21) (Table 3). The detected positives were significantly different for the presence of mutant and wild type DNA for both groups of dogs (Fig 5). To ensure an active infection is established in the infection control dogs and further to monitor the potential dynamics of wild-type *E. canis* infection, we performed qRT-PCR targeting 16S rRNA on the RNA samples recovered several days post wild-type *E. canis* infection challenge were assessed in all dogs, and RNA copies were estimated as in [66] (Fig 6 and Table S2). The data revealed the presence of significantly lower 16S rRNA copies in the Group 1 group throughout the assessment period (data shown in Table S2), whereas the infection control group dogs (Group 2) receiving wild-type *E. canis* infection had a rapid spike in the 16S rRNA copies, which peaked around day 14 post-infection challenge and declined to low levels there after which remained persisted till the study end point of assessment (Table S2). Animal body temperatures and weights remained similar for both groups throughout the study.

**Figure 5.**
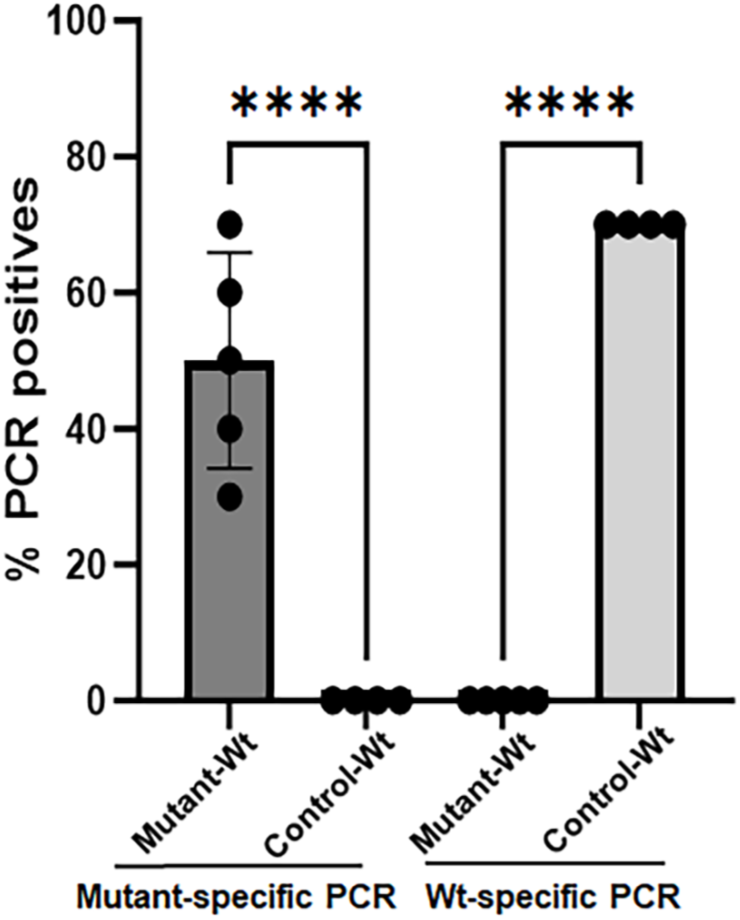
Vaccination with MLAV prevented the establishment of wild-type infection, as confirmed by PCR and culture recovery. Mutant- and wild-type-specific PCR analysis were performed on genomic DNAs extracted from blood samples collected at different time points following mutant infection and subsequent wild-type infection (Mutant-Wt), as well as after only wild-type infection (Control-Wt). The percentage of positives was used to determine differences in systemic bacterial presence between the groups. Mutant-specific PCR showing detection of the mutant DNA after wild-type infection only in the Mutant-Wt group (P < 0.0001 ****). Wild-type (Wt)-specific PCR detected wild-type-specific positives only in the blood DNAs of the control group (Control-Wt) (P < 0.0001 ****). Statistical differences between groups were determined using the one-way ANOVA with multiple comparisons.

**Figure 6.**
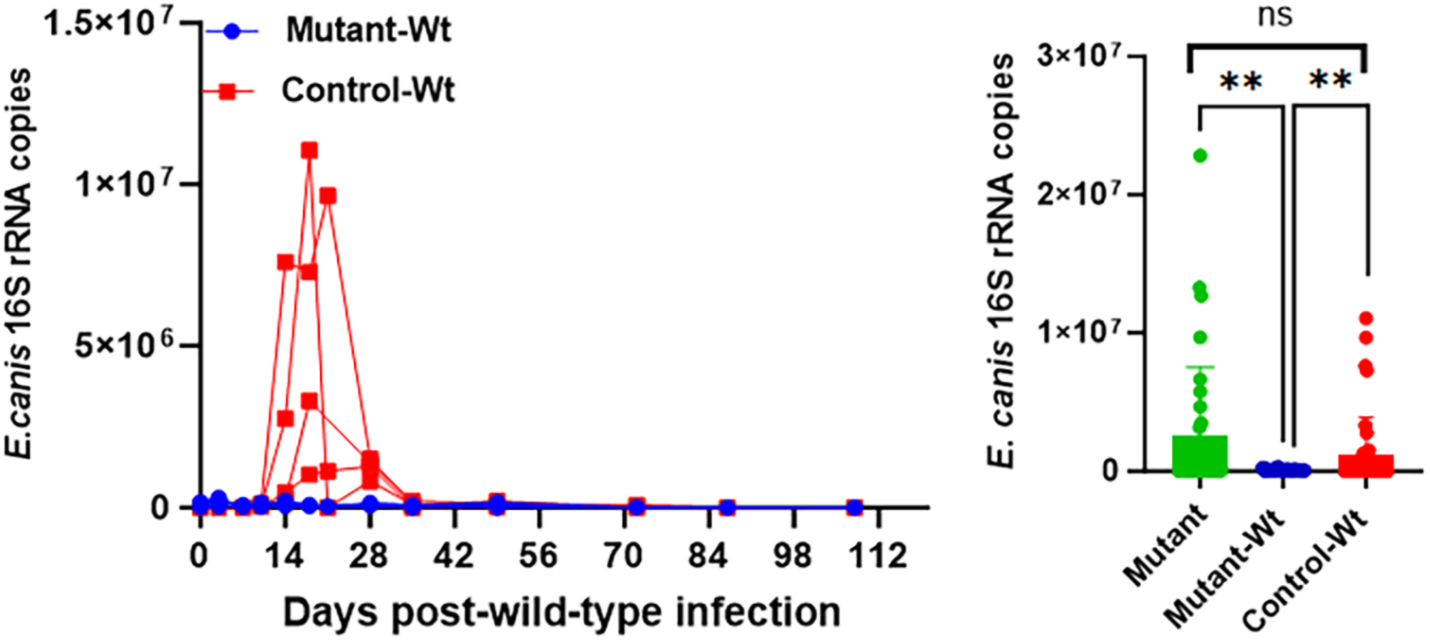
*E. canis 16s rRNA* copy numbers in blood-derived RNA assessed by TaqMan probe-based RT-PCR (qRT-PCR) following wild-type *E. canis* infection. Total RNA isolated from blood samples collected at various time points following wild-type infection from both groups of dogs was assessed by qRT-qPCR to quantify *E. canis* small subunit ribosomal RNA copies per 25 μl of blood (panel A). The RNA copies were significantly less in the prior mutant-infected group of dogs following the wild-type infection challenge compared with those observed following wild-type infection alone (*P* = 0.0065 **) or compared with those observed after the mutant infection only (described in Fig 2) (*P* = 0.0090 **) (panel B). Statistical significance was measured using an unpaired t-test with Welch’s correction.

**Table 3.** *E. canis* infection assessment by PCR targeting to map the presence of wild-type DNA (Wt-specific PCR).

|  | Dog ID | Days post-wildtype infection challenge* |  |  |  |  |  |  |  |  |  | Post-Doxycycline** |  |
| --- | --- | --- | --- | --- | --- | --- | --- | --- | --- | --- | --- | --- | --- |
|  |  | 3 | 7 | 10 | 14 | 18 | 21 | 28 | 35 | 49 | 72 | 11 | 32 |
| Mutant infected (Group 1) | AHL (M) | - | - | - | - | - | - | - | - | - | - | - | - |
|  | BAK (F) | - | - | - | - | - | - | - | - | - | - | - | - |
|  | BGK (F) | - | - | - | - | - | - | - | - | - | - | - | - |
|  | CUL (M) | - | - | - | - | - | - | - | - | - | - | - | - |
|  | DWL (M) | - | - | - | - | - | - | - | - | - | - | - | - |
| Controls (Group 2) | AVK (F) | - | - | + | + / C | + | + / C | + | + | - | + | - | - |
|  | BCK (F) | - | - | + | + / C | + | + / C | + | + | - | + | - | - |
|  | ECL (M) | - | - | + | + / C | + | + / C | + | + | - | + | - | - |
|  | GVL (M) | - | - | + | + / C | + | + / C | + | + | - | + | - | - |

\* + and – refer to PCR positives and negatives, respectively, and C refers to culture positives. M and F refer to male and female dogs, respectively.
\*\*All dogs in these two groups were treated with doxycycline for a month after 72 days of study assessment and then assessed for the presence of infection by PCR at the two time points listed.

#### 3.3.2. The CBC analysis showed an increase in the number of RBC only in the MLAV group dogs after wild-type infection challenge

We performed the CBC analysis following the subsequent wild-type infection challenge and compared the values to determine any differences between the two groups (Fig 7). Hematological abnormalities caused by *E. canis* infection, such as thrombocytopenia, may take months to recover, even with early clinical improvements [67, 68]. Platelet and WBC levels remained low in the mutant-infected group also following wild-type *E. canis* infection. However, the RBC levels improved significantly from day 10 onwards compared to the control group receiving only wild-type *E. canis* infection.

**Figure 7.**
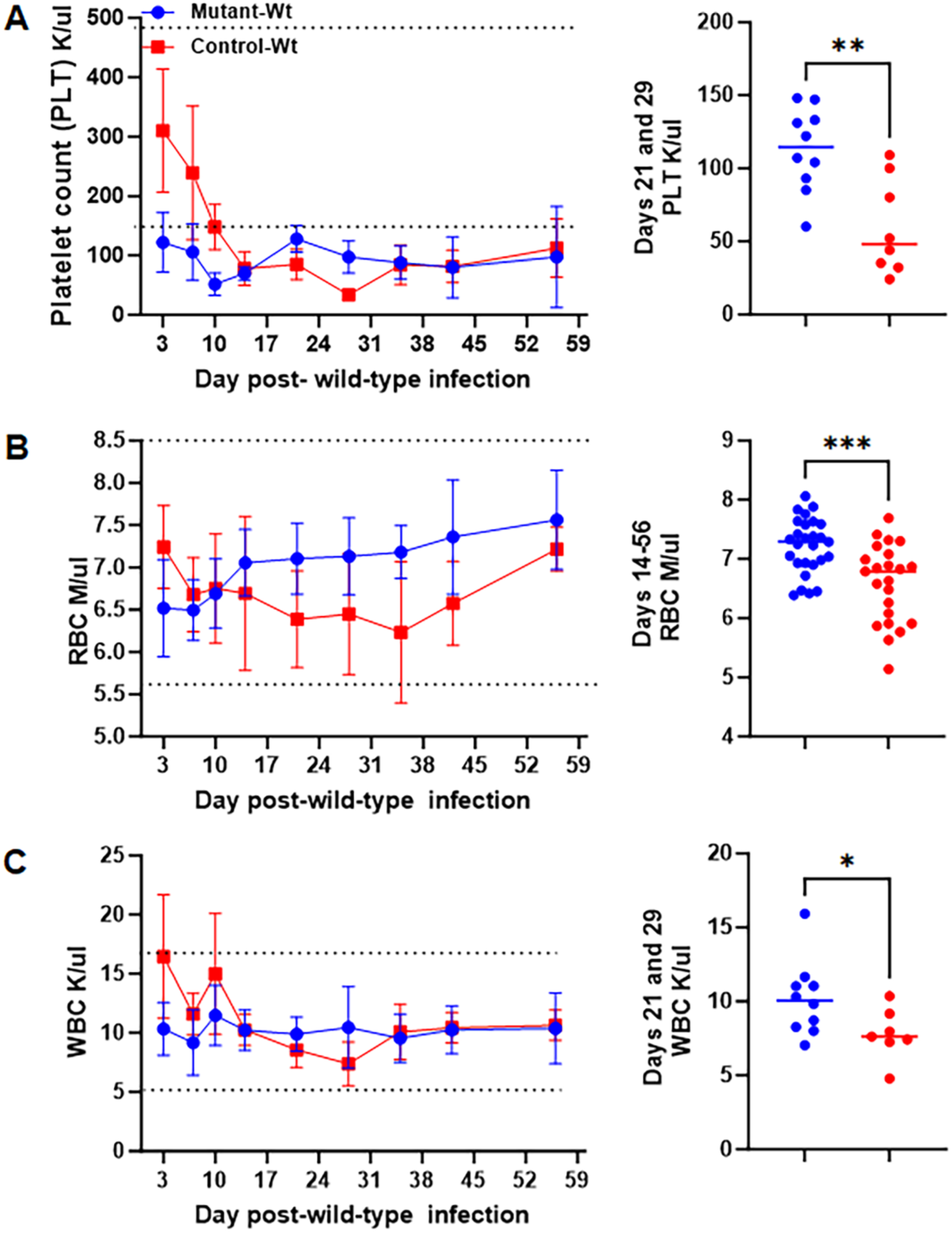
The CBC assessment following the wild-type *E. canis* infection challenge in naïve dogs or in dogs previously infected with the mutant. Both groups of dogs had a platelet drop (panel A). RBC counts significantly improved in dogs that had received prior mutant infection (P = 0.0335 *) (panel B). WBC counts were lower in naïve dogs receiving wild-type *E. canis* infection on days 21 and 28 post-infection compared with those previously infected with the mutant (panel C). Dotted lines refer to normal canine reference values. Statistical analysis was performed using an unpaired t-test with Welch’s correction for values from day 10 to 28 when the declines were evident after wild-type infection.

#### 3.3.3. Challenge with wild-type E. canis elicited an IgG response in all dogs

Immune response was measured by ELISA to detect antibodies against *E. canis* specific IgG over the course of the study. The presence of *E. canis-*specific IgG antibodies was evident for both groups of dogs following wild-type *E. canis* infection (Fig 8). The IgG response was similar for the mutant group following wild-type infection compared to the control group receiving only wild-type *E. canis* infection (Fig 8).

**Figure 8.**
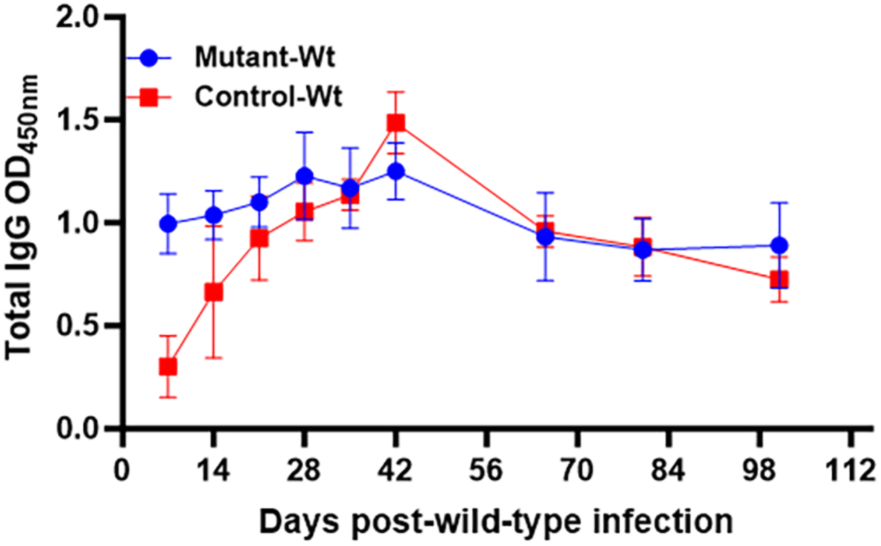
Total IgG response to *E. canis* antigens in dogs measured by ELISA. Plasma samples collected weekly following wild-type *E. canis* infection in the MLAV group and the control group were assessed by ELISA. IgG response following wild-type *E. canis* infection was similar in both groups.

#### 3.3.4. The MLAV and subsequent wild-type infection stimulated the specific cytokine responses

The panels of 11 cytokines were analyzed to determine the plasma cytokine profile following the mutant infection and after the wild-type infection challenge in the prior mutant-infected group of dogs, as well as for the wild-type infection controls (Fig 9). Animals from both groups showed detectible changes in four of the 11 cytokines assessed. Interleukin-8 (IL-8), also known as C-X-C motif chemokine ligand 8 (CXCL8), is a chemokine that recruits and activates neutrophils at inflammation sites during bacterial infections [32, 69–71]. Changes in IL-8 secretion levels of dogs post mutant infection did not significantly differ but decreased over the course of 21 days from their baseline (day 0) levels (Fig 9A). Whereas, independent of prior mutant infection or not, wild-type infection resulted in a significant rise in IL-8 starting on days 8 and 14 post infection and its levels in the prior mutant-infected dogs resulted in a significant drop on day 21 after wild-type infection. IL-12p40 is produced by immune cells, such as macrophages and dendritic cells, during intracellular bacterial infections, leading to the differentiation of naive T cells into type 1 helper T cells and stimulating the production of cytokines, including interferon gamma (IFN-γ), which help eliminate bacteria [71–75]. In contrast to IL-8, plasma IL-12p40 concentration steadily increased over time in dogs receiving the mutant as well as for the control dogs receiving the wild-type infection for the 21 days of assessment (Fig 9B). Prior mutant infection, however, caused in the IL-12p40 decline when this group of dogs received wild-type *E. canis* infection (Fig 9B). The stem cell factor (SCF) plays a significant role in hematopoiesis and is involved in stimulating the production of new peripheral blood cells (WBCs, RBCs, and platelets) during infection and after acute blood loss [76, 77]. Prior infection with the mutant caused no change in the SCF production and prevented its rise following wild-type *E. canis* infection in the mutant infection group dogs (Fig 9C). Contrary to this, its levels were significantly elevated in wild-type *E. canis* infection control dogs on days 14 and 21. IFN-γ is a major part of the TH1 response which is classically reported as the protective immune mechanism/response responsible for managing and eliminating infections [71, 73, 74, 78]. IFN-γ was significantly higher only after the mutant infection on day 7 and maintained higher concentration trends on days 14 and 21 (Fig 9D). Independent of the prior infection with the mutant or not, IFN-γ levels remained unchanged during the 21-day assessment period following wild-type *E. canis* infection challenge (Fig 9D). IFN-γ production was further assessed in PBMCs recovered from dogs and *in vitro* stimulated with *E. canis* antigens (Fig 10). We observed a trend towards higher numbers of PBMC producing IFN-γ on day 7 when measured by ELISPOT (Fig 10A) or higher IFN-γ recall responses by ELISA (Fig 10B) assays, independent of post-mutant infection or following wild-type *E. canis* infection challenge for this group and similarly for the wild-type infection control group dogs.

**Figure 9.**
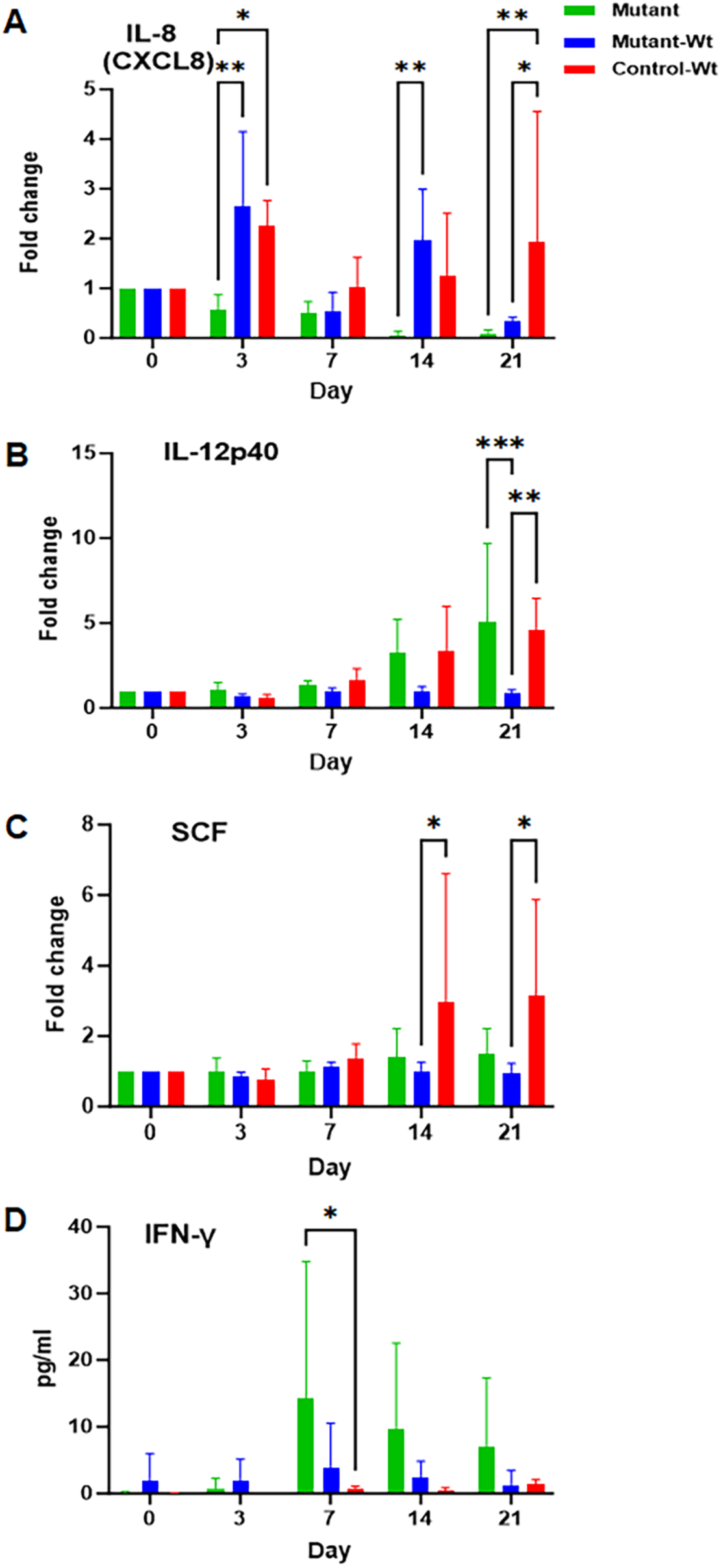
Plasma cytokine levels were measured using a Luminex bead-based assay. Cytokines were assessed following infection with the mutant (Mutant), or wild-type infection in naïve dogs (Wt), or wild-type infection following the prior mutant infection (Mutant-Wt). A) Notable changes in IL-8 levels were observed among the groups at different time points after the wild-type infection challenge; IL-8 expression was significantly higher on days 3 and 14 post-wild-type *E. canis* infection, independent of prior mutant infection or not (*P*=0.0012 ** and *P*=0.0144 *, respectively). On day 21 post wild-type infection, mutant infection or wild-type infection following MLAV had significantly less IL-8 compared to the dogs in the wild-type infection alone group (P = 0.0239 * and P = 0.0067 **). B) IL-12p40 levels were significantly lower following the mutant infection group dogs compared to dogs receiving wild-type *E. canis* infection alone or following the mutant infection on day 21 (*P* = 0.0003 *** and *P* = 0.0027 **, respectively). C) SCF levels were significantly higher only in the dogs receiving wild-type *E. canis* infection on days 14 and 21 post-infection (*P* = 0.0151 * and *P* = 0.0317 *, respectively. D) IFN-γ levels were significantly elevated on day 7 following infection with only the mutant strain (*P*= 0.0322 *), but not after wild-type *E. canis* infection, independent of prior infection with the mutant or not. Statistical significance was measured using a two-way ANOVA.

**Figure 10.**
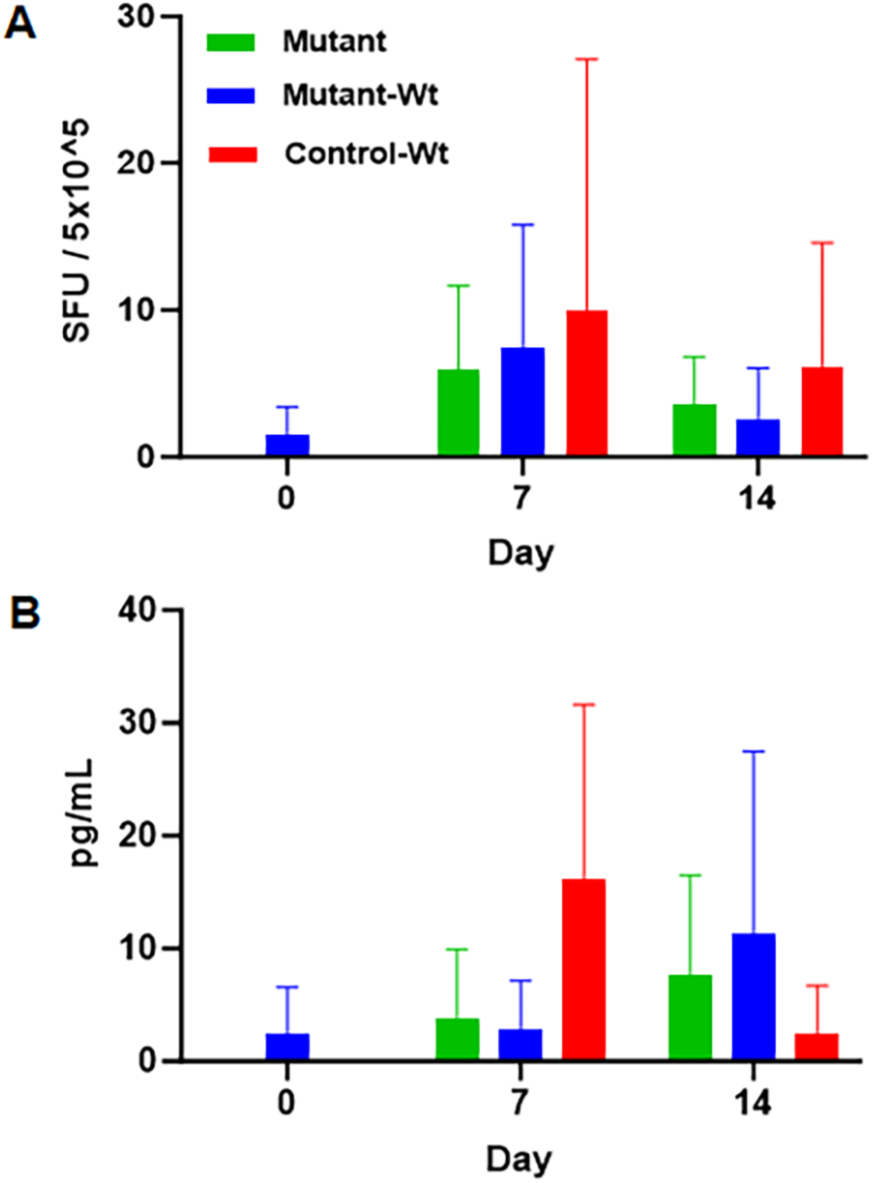
IFN-γ production in peripheral blood mononuclear cells (PBMC) collected over time. PBMCs were stimulated with *E. canis* antigens following infection with the mutant (Mutant), or wild-type infection in naïve dogs (Control-Wt), or wild-type infection following the prior mutant infection (Mutant-Wt). Groups 1 and 2 animals receiving wild-type infection alone or after mutant infection or following mutant infection alone induced IFN-γ production as measured by ELISPOT assay (panel A) or by ELISA (panel B). SFU indicating Spot Forming Unit.

#### 3.3.5. Treatment with doxycycline resulted in clearance of E. canis in circulation from all infected dogs

Ehrlichiosis is commonly treated with doxycycline [22, 44, 78], although some studies suggest that the treatment is ineffective in completely clearing the infection [46, 79]. After 72 days of infection monitoring, dogs in both Groups 1 and 2 were treated with doxycycline orally for one month. Subsequently, blood sampled from all dogs at two points post-treatment were tested by PCR to determine if the treatment was effective in clearing both mutant and wild-type *E. canis* (Tables 2 and 3). Neither the mutant nor the wild-type *E. canis* was detected in all dogs following doxycycline treatment by PCR, although we cannot rule out the possibility that the pathogen remains in dogs at levels below detection by PCR assays despite the treatment.

## 4. Discussion

Genetic modifications of obligate intracellular pathogens, such as those belong to the families Anaplasmataceae and Rickettsiaceae, serve as essential tools in understanding the molecular mechanisms important for pathogenesis and to identify virulence factors, as well as in developing methods of prevention. The ability to inactivate a gene of interest in these pathogens has remained challenging, likely due to the host-dependent nature and the bacterial extensive genome reductions [80, 81]. Despite the limitations, three targeted mutations in Rickettsiaceae family pathogens were reported; one each by an allelic exchange homologous recombination, a group II intron-based targeted mutagenesis, and a CRISPR-mediated mutagenesis are reported in three clinically important pathogenic *Rickettsia* species: *R. prowazekii* [82]*, R. rickettsii* [15], and *R. parkeri* [83]. Similarly, we reported targeted mutations in two Anaplasmataceae family pathogens: *A. marginale* and *E. chaffeensis* [11, 12, 47]. Using allelic exchange methods, we generated several targeted mutations in *E. chaffeensis*. The loss of function mutations in the *phtcp* genes of *E. chaffeensis* and *A. marginale* resulted in growth attenuation, suggesting that the gene is necessary for nutrient scavenging and survival. Indeed, we reported earlier that functional *phtcp* is required for metal ion homeostasis in *E. chaffeensis* [54]. As the *phtcp* is conserved within the Anaplasmataceae family bacteria [12, 54], we inactivated its gene ortholog from *E. canis* and assessed the mutant as a modified live attenuated vaccine (MLAV). Our previous targeted mutagenesis involved the insertion of an antibiotic resistance gene for a mutant generation. The present study extended the mutant generation with eliminating the need to include an antibiotic resistance gene. The mutated *E. canis* contained only the marker gene expressing mCherry protein, for which serial dilution coupled with gel agar-based plaque purification of the mutant provided an antibiotic-free mutant of *E. canis*. Eliminating the introduction of an antibiotic resistance cassette is valuable for a vaccine’s application, as it reduces the risk of antibiotic resistance development by the bacterial pathogens. Three independent experiments were performed to confirm the clonal purity and the presence of the mutation at the desired genomic location following which we tested and documented its value as a mutant modified live vaccine (supplementary Fig S2), along with the study flow chart (supplementary Fig S1) *E. canis* infection leading to CME is the most prevalent tick-borne disease of dogs globally [22, 24, 26]. The disease also gained significant US national attention in late 1960s as severe outbreaks among the US military dogs resulting from *E. canis* infections are reported [84]. Dogs recovering from the acute and subclinical disease remain positive for the pathogen lifelong and such dogs serve as reservoirs of infection facilitating *E. canis* life cycle maintained in nature via tick-transmission from the indoor *R. sanguineus sensu lato* (s.l.) complex [27, 28, 85]. The persistently infected dogs contribute to the risk of transmission to people from infected ticks. Human infections with *E. canis* are reported from Venezuela, Costa Rica, Mexico, Italy, and Montenegro [30–35]. Considering the global impact of this zoonotic pathogen, it is of critical importance to have a vaccine that protects the canine host which can also reduce zoonotic infections.

A prior study reported the description of cell culture-derived attenuated strain as having offered immune protection against virulent challenge [36]. It is unclear the molecular basis for the attenuation of *E. canis* resulting from the continuous cell culture passaging. Moreover, it is not uncommon for a cell culture attenuated rickettsial pathogen reverting to virulence [86]. With having no vaccines to prevent CME for nearly a century after its discovery and the lack of follow up investigations in defining the cell culture-attenuated vaccine described previously [36], the *phtcp* mutant serving as an MLAV offers a new path forward for advancing vaccine research. MLAVs are expected to offer stronger immune protection for intracellular bacterial pathogens as they have the capability to stimulate all arms of immunity: macrophage activation and the MHC II-mediated CD4 T cell activation leading to promoting the B cell activation and long-lasting memory responses [18, 87–90].

The *E. canis* mutant as an MLAV prevented the establishment of wild-type *E. canis* infection challenge. This is similar to prior evidence of immune protection reported with the *phtcp* mutations in *A. marginale* and *E. chaffeensis* infections [12, 17–21]. Whereas the wild-type infection in naïve dogs resulted in persistent systemic infection which was confirmed by PCR and culture. Nonetheless, no overt clinical signs such as fever were observed in naive dogs receiving wild-type *E. canis* infection. This likely reflects an establishment of a subclinical infection course under the controlled experimental conditions with intravenous infection challenge. Future investigations may require an infection challenge by tick transmission. The current study documenting persistent infection with the *E. canis* mutant in dogs is similar to prior documentation of persistent infection in cattle with the *phtcp* mutant of *A. marginale* which also protected wild-type pathogen infection challenges [12, 19]. Opposing to these observations, a similar *phtcp* mutant *E. chaffeensis* infection in the canine host results in both clearing the mutant as well as offering immune protection in preventing the wild-type infection progression [17, 18, 20, 21]. These data imply that, despite the close genetic similarities among the three rickettsial species, the pathogens have evolved unique abilities for establishing and maintaining infections in a host.

Although all hematological parameters were similar in dogs following infection with the mutant compared to wild-type *E. canis* infection, wild-type *E. canis* infection led to a marked increase in systemic IL-8 levels on days 3, 14, and 21 in the control group, suggesting a strong inflammatory response to wild-type infection, while after mutant infection, IL-8 levels significantly decreased. These results suggest that the immune response differs between infection with the mutant and wild-type *E. canis*. Conversely, subsequent wild-type infection of dogs previously infected with the mutant demonstrated a shorter period of increased IL-8 that was detected on days 3 and 14 post wild-type challenge. Nevertheless, systemic IL-12p40 secretion following infection was very similar between the mutant and wild-type *E. canis,* with a significant increase observed on day 21 post-infection. However, the levels remained significantly lower following wild-type infection in the prior mutant-infected group. This finding suggested that the primary systemic response to *E. canis* infection results in high systemic levels of IL-12p40; however, this robust systemic response is not seen with subsequent infection challenge, likely due to the prevention of the wild-type infection establishment. Interestingly, even though IL-12p40 is increased after both mutant and wild-type pathogen challenges, the downstream increase in IFN-γ is only seen after mutant infection, with detectable systemic IFN-γ in the blood of dogs on day 7 post-infection. This observation illustrates a clear biological difference between mutant and wild-type *E. canis* on the level of host-pathogen-immune interactions, suggesting that the *phtcp* deletion mutant did not block systemic IFN-γ production while wild-type infection inhibited its production. The SCF levels were significantly increased in the control group dogs on days 14 and 21 post wild-type infection compared to the dogs challenged with the mutant. While the mutant infection caused a decrease in WBC, RBC and platelet values similar to wild-type pathogen challenge, wild-type infection only animals had significantly higher systemic SCF during infection phase.

These data suggest that the blood cell concentrations were only maintained in the wild-type *E. canis* infected dogs due to a higher rate of cell replacement required to restore blood cell populations as indicated by increased SCF systemic levels. Dogs not previously exposed to the mutant as an MLAV had developed significant pancytopenia compared to MLAV dogs. These findings suggest that the mutant as a vaccine modulated the inflammatory cytokine milieu resulting in a more restrained systemic response which prevented the establishment of wild-type *E. canis* infection. T cell responses during intracellular bacterial infections play a predominant role in conferring protective immunity against infection than B cell responses [91, 92]. Consistent with this concept, we observed activation of T cell responses and evidence of T-cell memory, as indicated by increased IFN-γ production.

In summary, the data presented in the current study in creating the first mutation in *E. canis* and its application in assessing as an MLAV are novel. This study suggests that the targeted mutagenesis methods, which we initially developed for *E. chaffeensis*, are applicable to diverse Anaplasmataceae pathogens [11, 12, 47]. The molecular genetics development facilitated investigating a modified live vaccine in the physiologically relevant canine host, and it resulted in preventing the establishment of wild-type *E. canis* infection.

## Acknowledgements

This research was supported by the PHS grants #s R01AI152418 and R01AI070908 from the National Institute of Allergy and Infectious Diseases, NIH, USA.

## References

[1] Walker DH, Dumler JS. Emergence of the ehrlichioses as human health problems. Emerg Infect Dis. 1996;2:18–29.

[2] Paddock CD, Childs JE. *Ehrlichia chaffeensis*: a prototypical emerging pathogen. Clin Microbiol Rev. 2003;16:37–64.

[3] Bermúdez CSE, Troyo A. A review of the genus *Rickettsia* in Central America. Res Rep Trop Med. 2018;9:103–12.

[4] Walker DH, Paddock CD, Dumler JS. Emerging and re-emerging tick-transmitted rickettsial and ehrlichial infections. Medical Clinics of North America. 2008;92:1345–61.

[5] Uminski K, Kadkhoda K, Houston BL, Lopez A, MacKenzie LJ, Lindsay R, et al. Anaplasmosis: An emerging tick-borne disease of importance in Canada. IDCases. 2018;14:e00472.

[6] Kocan KM, de la Fuente J, Guglielmone AA, Melendez RD. Antigens and alternatives for control of *Anaplasma marginale* infection in cattle. Clin Microbiol Rev. 2003;16:698–712.

[7] Giraldo-Ríos C, Betancur O. Economic and health impact of the ticks in production animals. In: Abubakar M, Kanchana Perera P, editors. Ticks and Tick-Borne Pathogens. London: IntechOpen; 2018.

[8] Grasperge BJ, Wolfson W, Macaluso KR. Rickettsia parkeri infection in domestic dogs, Southern Louisiana, USA, 2011. Emerg Infect Dis. 2012;18:995–7.

[9] Afonso P, Lopes AP, Quintas H, Cardoso L, Coelho AC. *Ehrlichia canis* and *Rickettsia conorii* infections in shelter dogs: Seropositivity and implications for public health. Pathogens. 2024;13.

[10] McClure EE, Chavez ASO, Shaw DK, Carlyon JA, Ganta RR, Noh SM, et al. Engineering of obligate intracellular bacteria: progress, challenges and paradigms. Nat Rev Microbiol. 2017;15:544–58.

[11] Wang Y, Wei L, Liu H, Cheng C, Ganta RR. A genetic system for targeted mutations to disrupt and restore genes in the obligate bacterium, *Ehrlichia chaffeensis*. Sci Rep. 2017;7:15801.

[12] Hove P, Madesh S, Nair A, Jaworski D, Liu H, Ferm J, et al. Targeted mutagenesis in *Anaplasma marginale* to define virulence and vaccine development against bovine anaplasmosis. PLOS Pathog. 2022;18:e1010540.

[13] Fisher DJ, Beare PA. Recent advances in genetic systems in obligate intracellular human-pathogenic bacteria. Front Cell Infect Microbiol. 2023;13:1202245.

[14] Binet R, Maurelli AT. Transformation and isolation of allelic exchange mutants of *Chlamydia psittaci* using recombinant DNA introduced by electroporation. Proc Natl Acad Sci U S A. 2009;106:292–7.

[15] Noriea NF, Clark TR, Hackstadt T. Targeted knockout of the *Rickettsia rickettsii* OmpA surface antigen does not diminish virulence in a mammalian model system. mBio. 2015;6.

[16] Mueller KE, Wolf K, Fields KA. Gene deletion by fluorescence-reported allelic exchange mutagenesis in *Chlamydia trachomatis*. mBio. 2016;7:e01817–15.

[17] Nair AD, Cheng C, Jaworski DC, Ganta S, Sanderson MW, Ganta RR. Attenuated mutants of *Ehrlichia chaffeensis* induce protection against wild-type infection challenge in the reservoir host and in an incidental host. Infect Immun. 2015;83:2827–35.

[18] McGill JL, Nair AD, Cheng C, Rusk RA, Jaworski DC, Ganta RR. Vaccination with an attenuated mutant of *Ehrlichia chaffeensis* induces pathogen-specific CD4+ T cell immunity and protection from tick-transmitted wild-type challenge in the canine host. PLoS One. 2016;11:e0148229.

[19] Ferm J, Jaworski DC, Stoll I, Kleinhenz MD, Kocan KM, Madesh S, et al. Genetically modified live vaccine offers protective immunity against wild-type *Anaplasma marginale* tick-transmission challenge. Vaccine. 2024;42:126069.

[20] Madesh S, McGill J, Jaworski DC, Ferm J, Liu H, Fitzwater S, et al. Long-term protective immunity against *Ehrlichia chaffeensis* infection induced by a genetically modified live vaccine. Vaccines (Basel). 2024;12.

[21] Madesh S, McGill J, Jaworski DC, Ferm J, Ferm D, Liu H, et al. Prolonged immune response to tick-borne *Ehrlichia chaffeensis* infection using a genetically modified live vaccine. Vaccine. 2025;48:126730.

[22] Aziz MU, Hussain S, Song B, Ghauri HN, Zeb J, Sparagano OA. Ehrlichiosis in dogs: A comprehensive review about the pathogen and its vectors with emphasis on South and East Asian Countries. Vet Sci. 2022;10.

[23] Balmori-de la Puente A, Rodriguez-Escolar I, Collado-Cuadrado M, Infante Gonzalez-Mohino E, Vieira Lista MC, Hernandez-Lambrano RE, et al. Transmission risk of vector-borne bacterial diseases (*Anaplasma spp*. and *Ehrlichia canis*) in Spain and Portugal. BMC Vet Res. 2024;20:526.

[24] Beall MJ, Alleman AR, Breitschwerdt EB, Cohn LA, Couto CG, Dryden MW, et al. Seroprevalence of *Ehrlichia canis*, *Ehrlichia chaffeensis* and *Ehrlichia ewingii* in dogs in North America. Parasit Vectors. 2012;5:29.

[25] Bordim SC, Souza PM, Morales MJA, Fajardo HV, Santos HA, Rossi MF, et al. Molecular characterization of *Ehrlichia canis* in dogs from Brazil: a worldwide perspective. Vet Parasitol Reg Stud Reports. 2025;60:101243.

[26] Chakraborty A, Rath PK, Panda SK, Mishra BP, Dehuri M, Biswal S, et al. Molecular confirmation, epidemiology, and pathophysiology of *Ehrlichia canis* prevalence in Eastern India. Pathogens. 2024;13.

[27] Dantas-Torres F. Biology and ecology of the brown dog tick, *Rhipicephalus sanguineus*. Parasit Vectors. 2010;3:26.

[28] Gray J, Dantas-Torres F, Estrada-Pena A, Levin M. Systematics and ecology of the brown dog tick, *Rhipicephalus sanguineus*. Ticks Tick Borne Dis. 2013;4:171–80.

[29] Pascoe EL, Nava S, Labruna MB, Paddock CD, Levin ML, Marcantonio M, et al. Predicting the northward expansion of tropical lineage *Rhipicephalus sanguineus sensu lato* ticks in the United States and its implications for medical and veterinary health. PLOS ONE. 2022;17:e0271683.

[30] Perez M, Bodor M, Zhang C, Xiong Q, Rikihisa Y. Human infection with *Ehrlichia canis* accompanied by clinical signs in Venezuela. Ann N Y Acad Sci. 2006;1078:110–7.

[31] Perez M, Rikihisa Y, Wen B. *Ehrlichia canis*-like agent isolated from a man in Venezuela: antigenic and genetic characterization. J Clin Microbiol. 1996;34:2133–9.

[32] Garcia-Rosales L, Escarcega-Avila A, Ramirez-Lopez M, Manzanera-Ornelas D, Guevara-Macias E, Vaquera-Arteaga M, et al. Immune monitoring of paediatric patients infected with *Rickettsia rickettsii*, *Ehrlichia canis* and coinfected. Pathogens. 2022;11.

[33] Sgroi G, D’Alessio N, Veneziano V, Rofrano G, Fusco G, Carbonara M, et al. Ehrlichia canis in human and tick, Italy, 2023. Emerg Infect Dis. 2024;30:2651–4.

[34] Andrić B. Diagnostic evaluation of *Ehrlichia canis* human infections. Open Journal of Medical Microbiology. 2014;4.

[35] Bouza-Mora L, Dolz G, Solorzano-Morales A, Romero-Zuniga JJ, Salazar-Sanchez L, Labruna MB, et al. Novel genotype of *Ehrlichia canis* detected in samples of human blood bank donors in Costa Rica. Ticks Tick Borne Dis. 2017;8:36–40.

[36] Rudoler N, Baneth G, Eyal O, van Straten M, Harrus S. Evaluation of an attenuated strain of *Ehrlichia canis* as a vaccine for canine monocytic ehrlichiosis. Vaccine. 2012;31:226–33.

[37] Alves-Ribeiro BS, Duarte RB, Assis-Silva ZM, Gomes APC, Silva YA, Fernandes-Silva L, et al. *Ehrlichia canis* vaccine development: Challenges and advances. Vet Sci. 2024;11.

[38] Harrus S, Waner T. Diagnosis of canine monocytotropic ehrlichiosis (*Ehrlichia canis*): an overview. Vet J. 2011;187:292–6.

[39] Harrus S, Waner T, Bark H, Jongejan F, Cornelissen AW. Recent advances in determining the pathogenesis of canine monocytic ehrlichiosis. J Clin Microbiol. 1999;37:2745–9.

[40] Mylonakis ME, Koutinas AF, Breitschwerdt EB, Hegarty BC, Billinis CD, Leontides LS, et al. Chronic canine ehrlichiosis *(Ehrlichia canis)*: A retrospective study of 19 natural cases. Journal of the American Animal Hospital Association. 2004;40:174–84.

[41] Mylonakis ME, Ceron JJ, Leontides L, Siarkou VI, Martinez S, Tvarijonaviciute A, et al. Serum acute phase proteins as clinical phase indicators and outcome predictors in naturally occurring canine monocytic ehrlichiosis. Journal of Veterinary Internal Medicine. 2011;25:811–7.

[42] Bartsch RC, Greene RT. Post-therapy antibody titers in dogs with ehrlichiosis: follow-up study on 68 patients treated primarily with tetracycline and/or doxycycline. J Vet Intern Med. 1996;10:271–4.

[43] Breitschwerdt EB, Hegarty BC, Hancock SI. Doxycycline hyclate treatment of experimental canine ehrlichiosis followed by challenge inoculation with two *Ehrlichia canis* strains. Antimicrob Agents Chemother. 1998;42:362–8.

[44] Fourie JJ, Horak I, Crafford D, Erasmus HL, Botha OJ. The efficacy of a generic doxycycline tablet in the treatment of canine monocytic ehrlichiosis. J S Afr Vet Assoc. 2015;86:1193.

[45] Harrus S, Waner T, Aizenberg I, Bark H. Therapeutic effect of doxycycline in experimental subclinical canine monocytic ehrlichiosis: evaluation of a 6-week course. J Clin Microbiol. 1998;36:2140–2.

[46] Iqbal Z, Rikihisa Y. Reisolation of *Ehrlichia canis* from blood and tissues of dogs after doxycycline treatment. J Clin Microbiol. 1994;32:1644–9.

[47] Cheng C, Nair AD, Indukuri VV, Gong S, Felsheim RF, Jaworski D, et al. Targeted and random mutagenesis of *Ehrlichia chaffeensis* for the identification of genes required for *in vivo* infection. PLoS Pathog. 2013;9:e1003171.

[48] Gundelly P, Ransom E, Stewart Z, Ruch B, Jittirat A, Denny L, et al. Transmission of *Ehrlichia chaffeensis* from an organ donor to a kidney-pancreas transplant recipient. Transpl Infect Dis. 2025:e70107.

[49] Paddock CD, Folk SM, Shore GM, Machado LJ, Huycke MM, Slater LN, et al. Infections with *Ehrlichia chaffeensis* and *Ehrlichia ewingii* in persons coinfected with human immunodeficiency virus. Clin Infect Dis. 2001;33:1586–94.

[50] Paddock CD, Sumner JW, Shore GM, Bartley DC, Elie RC, McQuade JG, et al. Isolation and characterization of *Ehrlichia chaffeensis* strains from patients with fatal ehrlichiosis. J Clin Microbiol. 1997;35:2496–502.

[51] Scolarici MJ, Kuehler D, Osborn R, Doyle A, Schiffman EK, Garvin A, et al. Donor-derived ehrlichiosis caused by *Ehrlichia chaffeensis* from living donor kidney transplant. Emerg Infect Dis. 2025;31:587–90.

[52] Zhang H, Chang Z, Mehmood K, Wang Y, Rehman MU, Nabi F, et al. First report of *Ehrlichia* infection in goats, China. Microbial Pathogenesis. 2017;110:275–8.

[53] Yu S, Modarelli J, Tomeček JM, French JT, Hilton C, Esteve-Gasent MD. Prevalence of common tick-borne pathogens in white-tailed deer and coyotes in south Texas. International Journal for Parasitology: Parasites and Wildlife. 2020;11:129–35.

[54] Torres-Escobar A, Juarez-Rodriguez MD, Ganta RR. Mutations in *Ehrlichia chaffeensis* genes ECH_0660 and ECH_0665 cause transcriptional changes in response to zinc or iron limitation. J Bacteriol. 2021;203:e0002721.

[55] Cheng C, Ganta RR. Laboratory maintenance of *Ehrlichia chaffeensis* and *Ehrlichia canis* and recovery of organisms for molecular biology and proteomics studies. Curr Protoc Microbiol. 2008;Chapter 3:Unit 3A 1.

[56] Wang Y, Nair ADS, Alhassan A, Jaworski DC, Liu H, Trinkl K, et al. Multiple *Ehrlichia chaffeensis* genes critical for its persistent infection in a vertebrate host are identified by random mutagenesis coupled with *in vivo* infection assessment. Infect Immun. 2020;88.

[57] Green MR, & Sambrook, J. Molecular cloning: A laboratory manual. Fourth ed: Cold Spring Harbor Laboratory Press; 2012.

[58] Wick RR, Judd LM, Gorrie CL, Holt KE. Completing bacterial genome assemblies with multiplex MinION sequencing. Microb Genom. 2017;3:e000132.

[59] Li H, Durbin R. Fast and accurate long-read alignment with Burrows-Wheeler transform. Bioinformatics. 2010;26:589–95.

[60] Danecek P, Bonfield JK, Liddle J, Marshall J, Ohan V, Pollard MO, et al. Twelve years of SAMtools and BCFtools. Gigascience. 2021;10.

[61] Kolmogorov M, Yuan J, Lin Y, Pevzner PA. Assembly of long, error-prone reads using repeat graphs. Nature Biotechnology. 2019;37:540–6.

[62] Bolger AM, Lohse M, Usadel B. Trimmomatic: a flexible trimmer for Illumina sequence data. Bioinformatics. 2014;30:2114–20.

[63] Langmead B, Salzberg SL. Fast gapped-read alignment with Bowtie 2. Nat Methods. 2012;9:357–9.

[64] Wick RR, Holt KE. Polypolish: Short-read polishing of long-read bacterial genome assemblies. PLoS Comput Biol. 2022;18:e1009802.

[65] Wei L, Liu H, Alizadeh K, Juarez-Rodriguez MD, Ganta RR. Functional characterization of multiple *Ehrlichia chaffeensis* sodium (cation)/proton antiporter genes involved in the bacterial pH homeostasis. Int J Mol Sci. 2021;22.

[66] Sirigireddy KR, Ganta RR. Multiplex detection of *Ehrlichia* and *Anaplasma* species pathogens in peripheral blood by real-time reverse transcriptase-polymerase chain reaction. J Mol Diagn. 2005;7:308–16.

[67] Waner T. Hematopathological changes in dogs infected with *Ehrlichia canis.* 2008.

[68] Jean-Sebastien Palerme D, MSc, DACVIM. Ehrlichiosis in Dogs. 2025.

[69] Pease JE, Sabroe I. The role of interleukin-8 and its receptors in inflammatory lung disease: implications for therapy. Am J Respir Med. 2002;1:19–25.

[70] Miura K, Matsuo J, Rahman MA, Kumagai Y, Li X, Rikihisa Y. Ehrlichia chaffeensis induces monocyte inflammatory responses through MyD88, ERK, and NF-kappaB but not through TRIF, interleukin-1 receptor 1 (IL-1R1)/IL-18R1, or toll-like receptors. Infect Immun. 2011;79:4947–56.

[71] Rikihisa Y. Mechanisms of obligatory intracellular infection with *Anaplasma phagocytophilum*. Clin Microbiol Rev. 2011;24:469–89.

[72] Jeong B, Pahan K. IL-12p40 monomer: A potential player in macrophage regulation. Immuno. 2024;4:77–90.

[73] Liu J, Cao S, Kim S, Chung EY, Homma Y, Guan X, et al. Interleukin-12: an update on its immunological activities, signaling and regulation of gene expression. Curr Immunol Rev. 2005;1:119–37.

[74] Zheng H, Ban Y, Wei F, Ma X. Regulation of interleukin-12 production in antigen-presenting cells. Adv Exp Med Biol. 2016;941:117–38.

[75] Cooper AM, Khader SA. IL-12p40: an inherently agonistic cytokine. Trends Immunol. 2007;28:33–8.

[76] Broudy VC. Stem cell factor and hematopoiesis. Blood. 1997;90:1345–64.

[77] Wang W, Zhang Y, Dettinger P, Reimann A, Kull T, Loeffler D, et al. Cytokine combinations for human blood stem cell expansion induce cell-type- and cytokine-specific signaling dynamics. Blood. 2021;138:847–57.

[78] Cardoso SP, Honorio-Franca AC, Franca DCH, Silva LPS, Fagundes-Triches DLG, Neves MCB, et al. Effects of doxycycline treatment on hematological parameters, viscosity, and cytokines in canine monocytic ehrlichiosis. Biology (Basel). 2023;12.

[79] Wen B, Rikihisa Y, Mott JM, Greene R, Kim HY, Zhi N, et al. Comparison of nested PCR with immunofluorescent-antibody assay for detection of *Ehrlichia canis* infection in dogs treated with doxycycline. J Clin Microbiol. 1997;35:1852–5.

[80] Riffaud CM, Rucks EA, Ouellette SP. Persistence of obligate intracellular pathogens: alternative strategies to overcome host-specific stresses. Front Cell Infect Microbiol. 2023;13:1185571.

[81] Blanc G, Ogata H, Robert C, Audic S, Suhre K, Vestris G, et al. Reductive genome evolution from the mother of *Rickettsia*. PLOS Genetics. 2007;3:e14.

[82] Driskell LO, Yu XJ, Zhang L, Liu Y, Popov VL, Walker DH, et al. Directed mutagenesis of the *Rickettsia prowazekii* pld gene encoding phospholipase D. Infect Immun. 2009;77:3244–8.

[83] McGinn J, Wen A, Edwards DL, Brinkley DM, Lamason RL. An expanded genetic toolkit for inducible expression and targeted gene silencing in *Rickettsia parkeri*. Journal of Bacteriology. 2024;206:e00091–24.

[84] Kelch WJ. The canine ehrlichiosis (tropical canine pancytopenia) epizootic in Vietnam and its implications for the veterinary care of military working dogs. Mil Med. 1984;149:327–31.

[85] Dantas-Torres F. The brown dog tick, Rhipicephalus sanguineus (Latreille, 1806) (Acari: Ixodidae): From taxonomy to control. Veterinary Parasitology. 2008;152:173–85.

[86] Alhassan A, Liu H, McGill J, Cerezo A, Jakkula L, Nair ADS, et al. *Rickettsia rickettsii* whole-cell antigens offer protection against rocky mountain spotted fever in the canine host. Infect Immun. 2019;87.

[87] Bhattacharya P, Dey R, Dagur PK, Kruhlak M, Ismail N, Debrabant A, et al. Genetically modified live attenuated *Leishmania donovani parasites* induce innate immunity through classical activation of macrophages that direct the th1 response in mice. Infect Immun. 2015;83:3800–15.

[88] Chirkova TV, Naykhin AN, Petukhova GD, Korenkov DA, Donina SA, Mironov AN, et al. Memory T-cell immune response in healthy young adults vaccinated with live attenuated *i*nfluenza A (H5N2) vaccine. Clin Vaccine Immunol. 2011;18:1710–8.

[89] Rosch JW, Iverson AR, Humann J, Mann B, Gao G, Vogel P, et al. A live-attenuated pneumococcal vaccine elicits CD4+ T-cell dependent class switching and provides serotype independent protection against acute otitis media. EMBO Mol Med. 2014;6:141–54.

[90] Altenburg AF, Kreijtz JH, de Vries RD, Song F, Fux R, Rimmelzwaan GF, et al. Modified vaccinia virus ankara (MVA) as production platform for vaccines against influenza and other viral respiratory diseases. Viruses. 2014;6:2735–61.

[91] Stabel JR. Transitions in immune responses to *Mycobacterium paratuberculosis*. Vet Microbiol. 2000;77:465–73.

[92] Vitry MA, De Trez C, Goriely S, Dumoutier L, Akira S, Ryffel B, et al. Crucial role of gamma interferon-producing CD4+ Th1 cells but dispensable function of CD8+ T cell, B cell, Th2, and Th17 responses in the control of Brucella melitensis infection in mice. Infect Immun. 2012;80:4271–80.

