## Supplementary figures and images for "Targeted mutagenesis in *Ehrlichia canis* deleting the phage head-to-tail connector protein gene and its assessment as a vaccine candidate preventing canine ehrlichiosis"

### Supplementary Fig 1 and 2

**Figure S1**

#### FLOW CHART OF THE RESEARCH OUTLINE

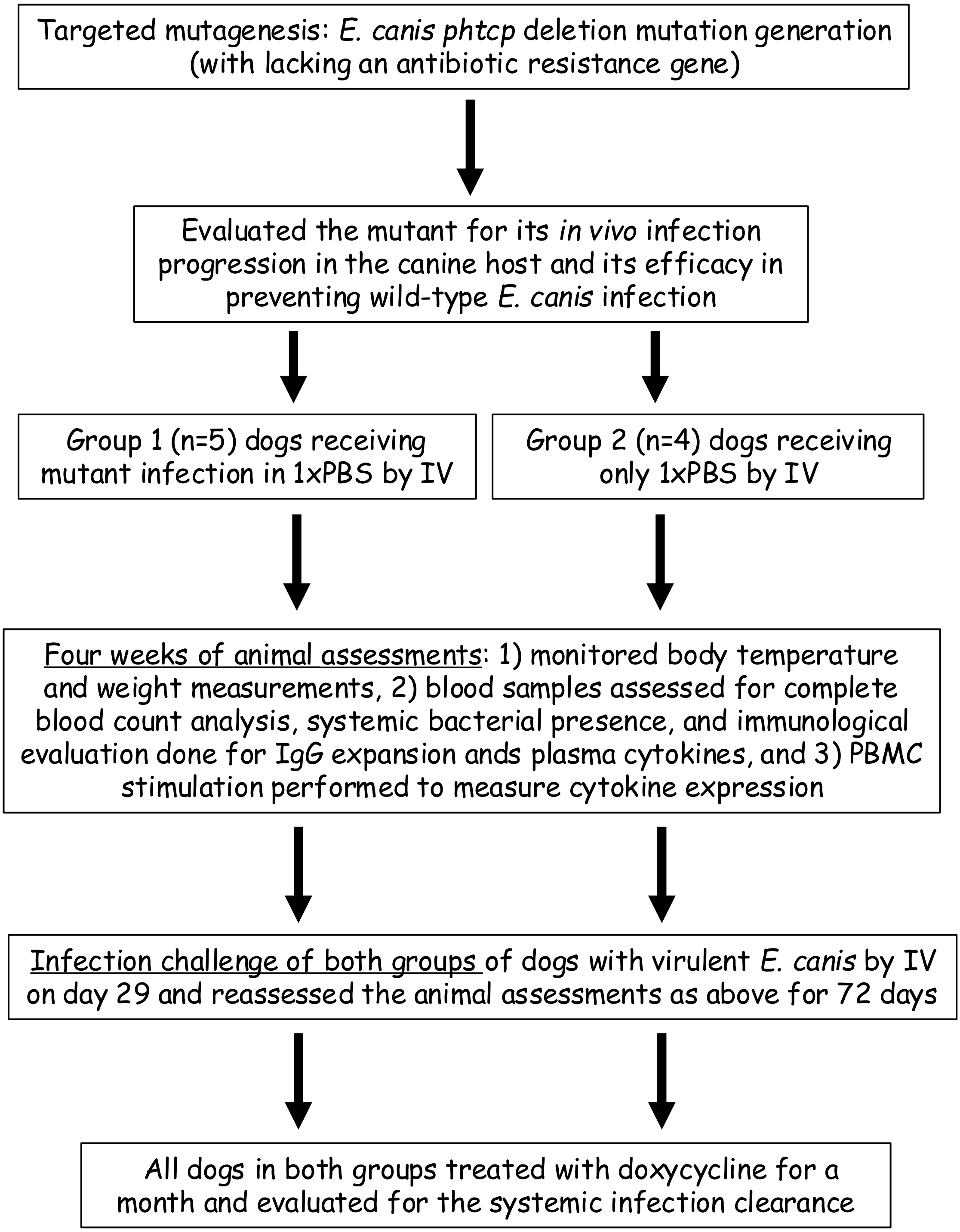

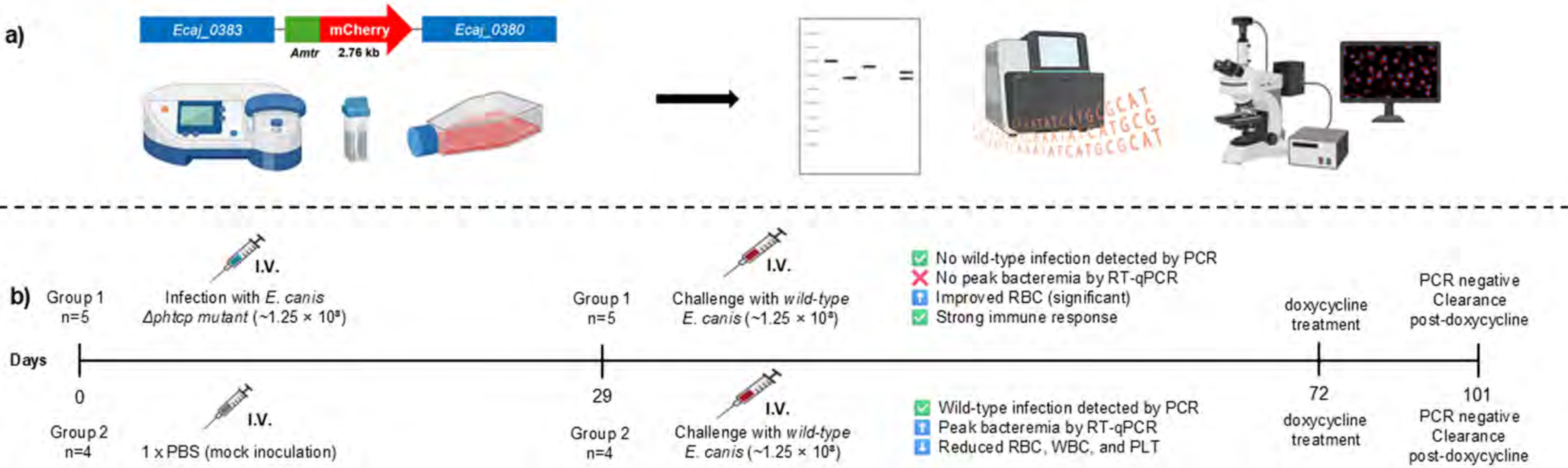
